# The Embryonic Origin of Primordial Germ Cells in the Tardigrade *Hypsibius exemplaris*

**DOI:** 10.1101/2023.01.02.522500

**Authors:** Kira L. Heikes, Mandy Game, Frank W. Smith, Bob Goldstein

**Author notes:** Corresponding author: Department of Biology, University of North Carolina at Chapel Hill, 616 Fordham Hall, Campus Box 3280, Chapel Hill, NC 27599-3280.

## Abstract

Primordial germ cells (PGCs) give rise to gametes – cells necessary for the propagation and fertility of diverse organisms. Current understanding of PGC development is limited to the small number of organisms whose PGCs have been identified and studied. Expanding the field to include little-studied taxa and emerging model organisms is important to understand the full breadth of the evolution of PGC development. In the phylum Tardigrada, no early cell lineages have been identified to date using molecular markers. This includes the PGC lineage. Here, we describe PGC development in the model tardigrade *Hypsibius exemplaris*. The four earliest-internalizing cells (EICs) exhibit PGC-like behavior and nuclear morphology. The location of the EICs is enriched for mRNAs of conserved PGC markers *wiwi1* (water bear *piwi* 1) and *vasa*. At early stages, both *wiwi1* and *vasa* mRNAs are detectable uniformly in embryos, which suggests that these mRNAs do not serve as localized determinants for PGC specification. Only later are *wiwi1* and *vasa* enriched in the EICs. Finally, we traced the cells that give rise to the four PGCs. Our results reveal the embryonic origin of the PGCs of *H. exemplaris* and provide the first molecular characterization of an early cell lineage in the tardigrade phylum. We anticipate that these observations will serve as a basis for characterizing the mechanisms of PGC development in this animal.

## Introduction

As cells funnel to their ultimate fates in development, one set of cells maintains a direct connection to the next generation. These are the primordial germ cells (PGCs), considered immortal because from PGCs arise gametes, the cellular basis of reproduction in many organisms (Kirkwood, 1987; Smelick and Ahmed, 2005; Strome and Updike, 2015). The proper specification of PGCs is therefore vital for the propagation of such organisms. Improper or ectopic PGC specification can lead to infertility, disorders in sex development, and an increased risk for germ cell cancer in humans (Hersmus et al., 2017). Despite the crucial nature of PGC development, current knowledge of PGC development is restricted to a small number of species. This is partly due to the limited number of organisms in which PGC lineages have been identified (Extavour and Akam, 2003; Whittle and Extavour, 2017). Identification of the embryonic origins of PGCs in a greater breadth of organisms is a key step toward building our understanding of the evolution of PGC development.

Throughout Metazoa, PGCs exhibit conserved features. These include internalization and migration to the future site of the gonad (Campanale et al., 2014; Dumstrei et al., 2004; Dzementsei and Pieler, 2014; Kunwar et al., 2006; Raz, 2004; Richardson and Lehmann, 2010; Sonnenblick, 1941). Additionally, PGCs typically exhibit cell cycle arrest for a period of time, until they are induced to proliferate in the gonad and differentiate into germ stem cells, which will ultimately propagate the gametes (Huettner, 1923; Pehrson and Cohen, 1986; Sekl et al., 2007; Sonnenblick, 1941; Su et al., 1998). This cell cycle arrest is important to protect the genome of PGCs from premature or improper differentiation (Strome and Updike, 2015). During the time of cell cycle arrest, PGCs often exhibit diffuse DNA staining, a reflection of a pluripotent cell state (Belew et al., 2021; D’Orazio et al., 2021; Lebedeva et al., 2018; Risley, 1933; Schaner et al., 2003; Yön and Akbulut, 2015). PGC fate is established and/or protected by a conserved set of genes; among these, the most widely used are the translational repressor Nanos, the nuclear argonaute protein Piwi, and the RNA helicase Vasa (Ewen-Campen et al., 2010; Juliano et al., 2010). Enriched presence of these genes’ products at the RNA and/or the protein level is often used as a marker of PGC fate (Chang et al., 2006; Ewen-Campen et al., 2013; Juliano et al., 2010; Leclère et al., 2012; Yoon et al., 1997).

Much is known about the PGC lineage in common model systems, such as the arthropod *Drosophila melanogaster* and the nematode *Caenorhabditis elegans* (Strome, 2005; Wang and Seydoux, 2013; Williamson and Lehmann, 1996). One phylum closely related to Arthropoda and Nematoda is Tardigrada (Aguinaldo et al., 1997). Members of this phylum are known for their ability to survive in the face of extreme conditions (Reviewed in Schill, 2018). Since the establishment of methods for culturing and studying the tardigrade species *Hypsibius exemplaris*, this species has grown as a model system for developmental, evolutionary, and extreme tolerance studies (Boothby, 2018; Gabriel et al., 2007; Goldstein, 2022b, 2022a, 2018; Heikes and Goldstein, 2018a; McGreevy et al., 2018; Mcnuff, 2018; Smith, 2018; Smith and Gabriel, 2018; Tenlen, 2018). Knowledge of the PGC lineage in the tardigrade phylum is limited to predictions made on the basis of light microscopy observations (Hejnol and Schnabel, 2006, 2005; Kaufmann, 1851; Marcus, 1929; von Erlanger, 1895; von Wenck, 1914). There is currently no definitive molecular evidence for which cells normally give rise to the PGCs, or any cell fate, in this phylum. Molecular studies thus far have uncovered various aspects of tardigrade embryonic development, such as development of the body plan, patterning and structure of the nervous system, and composition and expression patterns of the Wnt signaling family (Chavarria et al., 2021; Gabriel and Goldstein, 2007; Game and Smith, 2020; Smith et al., 2018, 2017, 2016; Smith and Goldstein, 2017; Smith and Jockusch, 2014). A molecular approach in tardigrades holds great promise in identifying embryonic cell lineages, like the PGCs.

In this study, we have characterized the PGC lineage in *H. exemplaris* through differential interference contrast (DIC) microscopy to ascertain cellular behaviors and cell position, chromogenic and fluorescent *in situ* hybridization (CISH and FISH, respectively) of conserved PGC markers *wiwi1* (water bear *piwi* 1) and *vasa* to molecularly distinguish the PGCs, and manual lineage tracing of the PGC precursor cells from 4D DIC videos to uncover the embryonic origin of the PGCs. The observations we report here support the hypothesis that the four EICs represent the PGC lineage of *H. exemplaris* embryos. Our results describe PGC development in *H. exemplaris* and provide the first molecular evidence for an early cell lineage in a tardigrade species.

## Results

### The four earliest-internalizing cells exhibit PGC-like behavior and chromatin morphology

We filmed *H. exemplaris* embryos by DIC microscopy (Video 1). Early embryonic development in *H. exemplaris* consists of many cell divisions in a compact ball of cells with no apparent blastocoel, as previously described (Gabriel et al., 2007). During this time, cells remain in a single layer, with nuclei positioned apically after each division and cell bodies extending towards the middle. Some cells briefly move into the middle of the embryo at various times during their cell cycles but return to the surface with apically-positioned nuclei during or after mitosis. No cells internalize and remain internalized until approximately 8 hours post laying (hpl), at which point a group of four cells are the first to internalize and remain inside, not returning to the surface. These four earliest-internalizing cells (EICs) were previously speculated to be germ cell precursors (Gabriel et al., 2007) based on observations by others who noted that cells in the position of the EICs in two species of tardigrades matched the position of the future germline (Hejnol and Schnabel, 2005; Marcus, 1929).

We observed that the four EICs underwent cell cycle arrest just before or after internalization. Cell divisions among the four EICs were not evident by DIC microscopy through at least the elongation stage; i.e. the four cells were seen to remain throughout this period (n = 9 embryos). The four EICs also migrated extensively once internalized (Figs. 1A-A”’, Video 1). As cell divisions continued and most of the cells in the embryo decreased in size as a result, the four EICs became increasingly apparent as four cells that remained large. After migrating, the four EICs were centrally located in the third trunk segment, near but distinct from the developing midgut (seen by the presence of birefringent granules under DIC light, evident around the segmentation stage, Fig. 1A’’’, D and E). This is the future location of the gonad in adults of this species, based on previous observations (Gross et al., 2019; Jezierska et al., 2021). Finally, the four EICs exhibited PGC-like chromatin morphology upon staining with DAPI: the DNA signal was visibly more diffuse in EICs than in surrounding cells (Figs. 1B and 1C; quantified in Supp. Fig. 1). This suggests that compared to surrounding, presumptively somatic cells, EIC cells have chromatin that is less compact. These characteristics of the four EICs led us to hypothesize that the four EICs are the PGCs of *H. exemplaris*.

**Figure 1.**
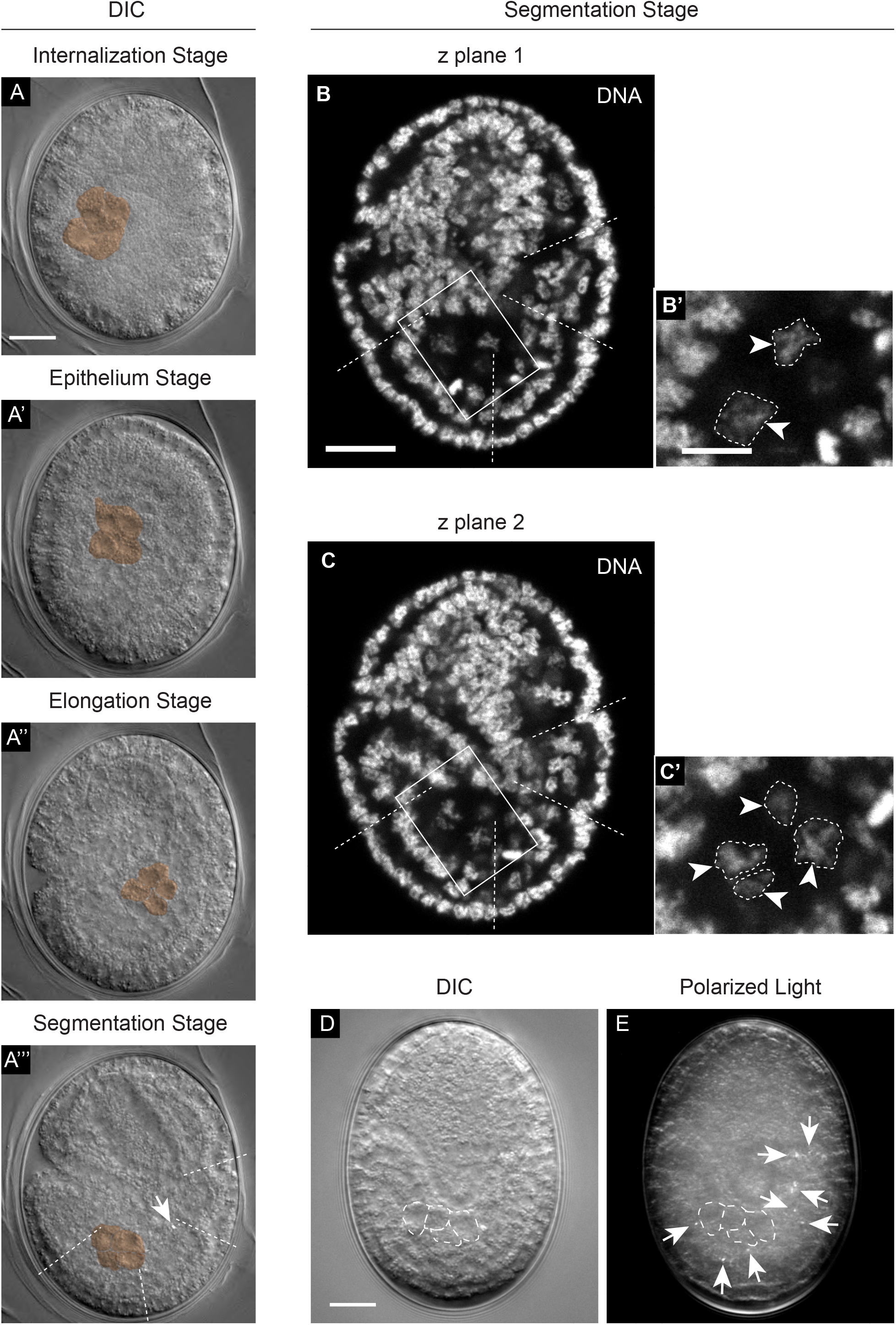
The four earliest-internalizing cells migrate to the future location of the gonad and exhibit PGC-like chromatin morphology. (A) DIC images showing the four EICs at four developmental stages (EICs manually colored with transparent orange overlay, scale bar = 10 μm) and their ultimate positioning in the third trunk segment near developing midgut, evident by the presence of birefringent granules (arrow indicating a birefringent granule in A’’’, approximate segment boundaries indicated by dotted lines). (B and C) DAPI-stained embryos at the segmentation stage showing the diffuse chromatin staining of cells in the position of the four EICs (each a single z plane, approximate segment boundaries indicated by dotted lines, scale bar = 10 μm). (B’ and C’) Expanded views of the boxed regions in B and C show the diffuse nature of EIC chromatin morphology (chromatin manually outlined with dotted lines and indicated by arrowheads, scale bar of enlarged region = 4 μm). Two of the four EIC nuclei are visible in the plane in B and B’, while all four EIC nuclei are visible in the slightly lower z plane in C and C’. (D and E) Position of the EICs at the segmentation stage relative to developing midgut, shown with an image of the four EICs by DIC light (D) and birefringent granules by polarized light (E), taken sequentially in a segmentation stage embryo (EICs outlined with white dotted lines and arrows indicating birefringent granules, scale bar = 10 μm).

### *wiwi1* mRNA is enriched in the region of the four earliest-internalizing cells

We sought to determine if molecular signatures of the PGC fate were present in the EICs. We were unable to identify a homolog of *nanos* in genomes or transcriptomes published for this system or for another tardigrade species, *Ramazzottius varieornatus* (Hashimoto et al., 2016; Levin et al., 2016; Yoshida et al., 2019, 2017). We identified and examined the enrichment of homologs of the conserved PGC markers *piwi* and *vasa*.

Using the *Drosophila melanogaster* Piwi protein sequence as a query, we identified two putative *H. exemplaris* homologs for Piwi by tBLASTn search and confirmed these as Piwi homologs by reciprocal BLASTp search. We named these *wiwi1* and *wiwi2* (water bear *piwi* 1 and 2). We performed a protein sequence alignment and generated a maximum likelihood phylogenetic reconstruction demonstrating a possible evolutionary relationship between Wiwi1 and 2 and other *H. exemplaris* argonaute homologs with known Piwi and argonaute homologs from other systems (Fig. 2 A). We also performed domain analyses and aligned the conserved PIWI and PAZ domains of Wiwi1 and 2 to their counterparts from homologs in other systems (Fig. 2 B and C). There was 30.31% protein sequence conservation between full-length Wiwi1 and *D. melanogaster* Piwi. There was 28.48% protein sequence conservation between full-length Wiwi2 and *D. melanogaster* Piwi.

**Figure 2.**
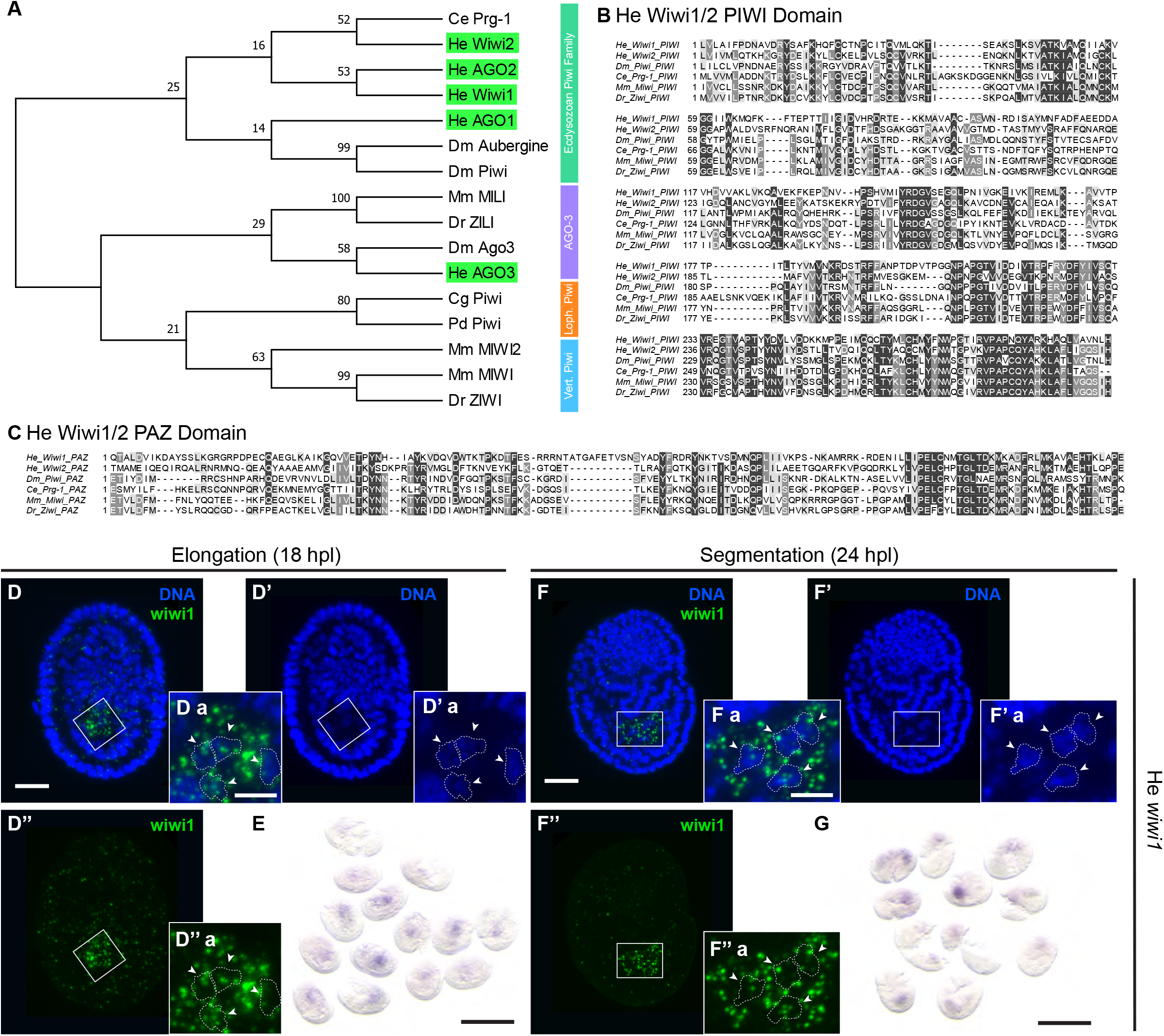
*wiwi1* mRNA surrounds the nuclei of the four large, undivided cells defined as the EICs. (A) Maximum-likelihood phylogenetic reconstruction of Piwi and related Argonaute amino acid sequences. Branch support out of 100 is given at each node. *H. exemplaris* sequences are highlighted in green. Argonaute protein families are indicated by colored bars to the right of the tree. Species name abbreviations are defined in Table 2. (B) Alignment of PIWI domain among Piwi homologs from several species. (C) Alignment of PAZ domain among Piwi homologs from several species. (D) Enrichment of *wiwi1* by FISH in elongation stage embryos (18 hpl, n = 3 experiments, 5, 6, and 9 embryos, respectively, scale bar = 10 μm). (F) Enrichment of *wiwi1* by FISH in segmentation stage embryos (24 hpl, n = 3 experiments, 4, 9, and 9 embryos, respectively, scale bar = 10 μm). Chromatin stained with DAPI is shown in blue, and *wiwi1* mRNA staining is shown in green. (Da – D’’’a and Fa – F’’’a) Enlargements of boxed regions in D and F show four nuclei (outlined manually and indicated with arrowheads) overlapping with the region enriched for *wiwi1* mRNAs (scale bar of enlarged region = 4 μm). (E) Widefield image of enrichment for *wiwi1* by CISH in elongation stage embryos (18 hpl, n = 2 experiments, 9 and 14 embryos, respectively, scale bar = 100 μm). (G) Widefield image of enrichment for *wiwi1* by CISH in segmentation stage embryos (24 hpl, n = 3 experiments, 10, 12 and 11 embryos, respectively, scale bar = 100 μm).

To see where *wiwi1* and *wiwi2* mRNAs were present in *H. exemplaris* embryos, we performed chromogenic in situ hybridization (CISH) in both elongation stage (18 hpl) and segmentation stage (24 hpl) embryos. There was a consistent, distinct enrichment of *wiwi1* signal in a small region on one end of the embryo at both 18 hpl and at 24 hpl (Fig. 2 E and G). Staining for *wiwi2* mRNAs did not yield consistent results (not shown). Therefore, we chose to use *wiwi1* as a consistent PGC marker going forward. To resolve exactly where the *wiwi1* signal enrichment was localized, we performed fluorescent in situ hybridization (FISH) with tyramide signal amplification. A set of cells toward the posterior, located in an internal layer of the third trunk segment at the segmentation stage, was enriched for *wiwi1* mRNAs (Fig. 2 D and F). The probe signal was punctate and present in the cytoplasm surrounding the nuclei of the four large, undivided cells that we defined above as the EICs (Fig. 1).

To understand how many cells were enriched for *wiwi1* mRNAs in this region of the embryo, we manually cropped the region around the enriched *wiwi1* signal in a segmentation stage embryo and produced a 3-D reconstruction of this region (using Imaris) and examined the number of nuclei engulfed by the *wiwi1* signal (Video 2). There were at least four nuclei surrounded by the enriched *wiwi1* signal. These four nuclei also exhibited the PGC-like (diffuse) chromatin morphology by DAPI staining, as described above. We conclude that the EICs are enriched for mRNAs of *wiwi1*, a conserved PGC marker.

### *vasa* mRNA is enriched in the region of the four earliest-internalizing cells

We repeated this molecular characterization with another conserved PGC marker, *vasa*. Using the *Drosophila melanogaster* Vasa protein sequence as a query, we identified one putative *H. exemplaris* homolog of Vasa by tBLASTn search and confirmed this by reciprocal BLASTp search. We performed a protein sequence alignment and generated a maximum likelihood phylogenetic reconstruction demonstrating a possible evolutionary relationship between *H. exemplaris* Vasa and Belle, another *H. exemplaris* RNA helicase homolog, with known homologs from other systems (Fig. 3 A). We also performed domain analysis and aligned the conserved c-terminal DEXD and Helicase domains of Vasa to their counterparts from other systems (Fig. 3 B and C). There was 46.67% protein sequence conservation between full-length *H. exemplaris* Vasa and *D. melanogaster* Vasa.

**Figure 3.**
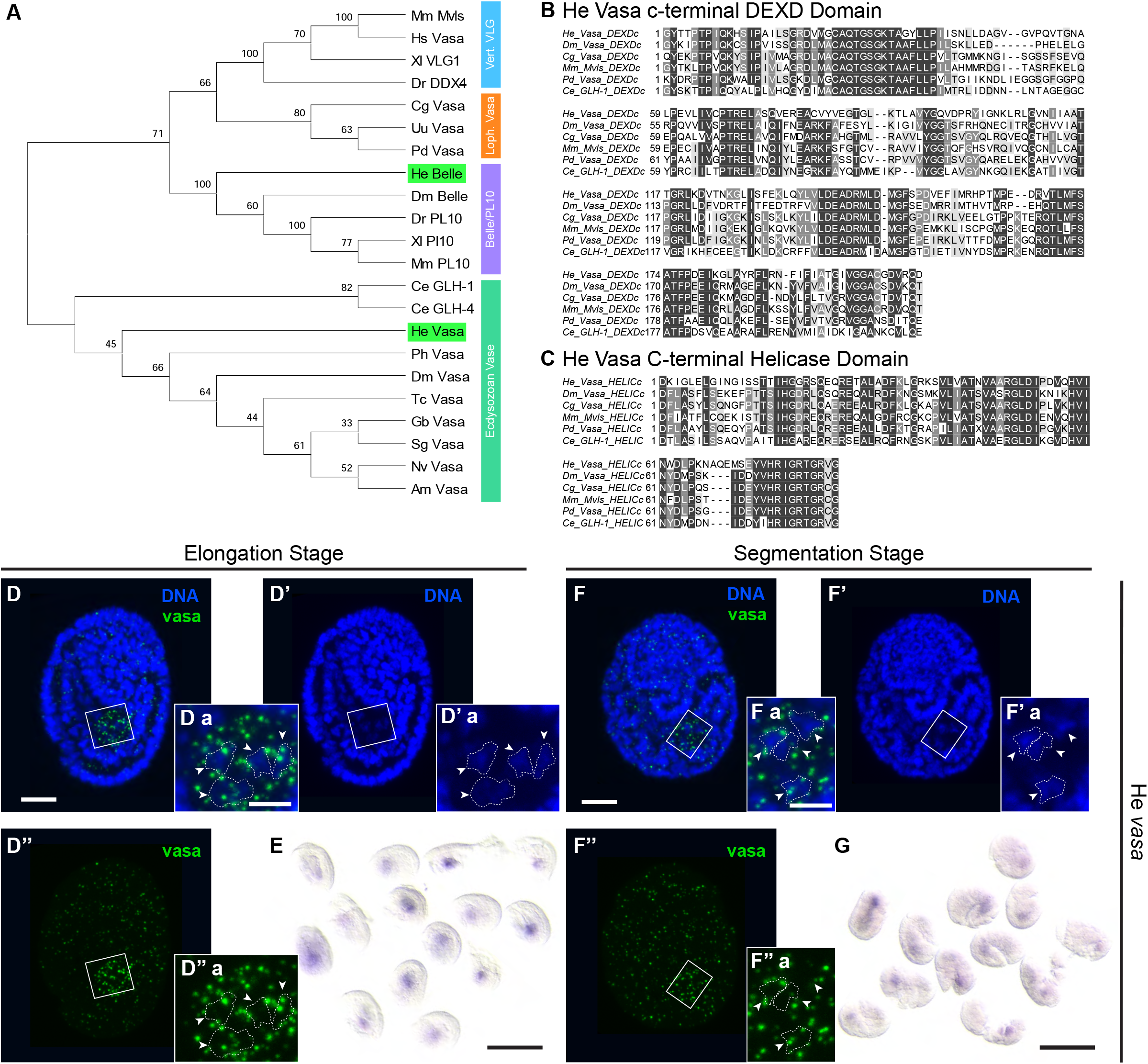
*vasa* mRNA surrounds the nuclei of the four large, undivided cells defined as the EICs. (A) Maximum-likelihood phylogenetic reconstruction of Vasa and related RNA helicase amino acid sequences. Branch support out of 100 is given at each node. *H. exemplaris* sequences are highlighted in green. RNA helicase protein families are indicated by colored bars to the right of the tree. Species name abbreviations are defined in Table 2. (B) Alignment of DEXD domain among Vasa homologs from several species. (C) Alignment of c-terminal Helicase domain among Vasa homologs from several species. (D-E) Elongation stage embryos (18 hpl). (F-G) Segmentation stage embryos (24 hpl). (D) Enrichment for *vasa* by FISH in elongation stage embryos (18 hpl, n = 3 experiments, 8, 9, and 9 embryos, respectively, scale bar = 10 μm). (F) Enrichment for *vasa* by FISH in segmentation stage embryos (24 hpl, n = 1 experiment, 9 embryos, scale bar = 10 μm). Chromatin stained with DAPI is shown in blue, and *vasa* mRNA staining is shown in green. (Da – D’’’a and Fa – F’’’a) Enlargements of boxed regions in D and F show four nuclei (outlined manually and indicated with arrowheads) overlapping with the region enriched for *vasa* mRNAs (scale bar of enlarged region = 4 μm). (E) Widefield image of enrichment for *vasa* by CISH in elongation stage embryos (18 hpl, n = 2 experiments, 9 and 14 embryos, respectively, scale bar = 100 μm). (G) Widefield image of enrichment for *vasa* by CISH in segmentation stage embryos (24 hpl, n = 2 experiments, 12 and 12 embryos, respectively, scale bar = 100 μm).

To see where *vasa* mRNAs were present, we performed chromogenic in situ hybridization (CISH) in both elongated (18 hpl) and segmented (24 hpl) embryos. Similar to *wiwi1*, there was a distinct enrichment of *vasa* signal in a small region on one end of the embryo (Fig. 3 E and G). To resolve exactly where this signal enrichment was localized, we performed fluorescent in situ hybridization (FISH) with tyramide signal amplification. As with *wiwi1*, a set of cells towards the posterior, located in an internal layer of the third trunk segment at the segmentation stage, was enriched for *vasa* mRNAs (Fig. 3 D and F). This probe signal likewise was punctate and present in the cytoplasm surrounding the nuclei of the four large, undivided cells that we defined above as the EICs.

As with *wiwi1*, to understand how many cells were enriched for *vasa* mRNAs in this region of the embryo, we manually cropped the region around the enriched *vasa* signal in an elongation stage embryo and produced a 3-D reconstruction of this region (using Imaris) and examined the number of nuclei engulfed by the *vasa* signal (Video 3). There were at least four nuclei overlapping with the enriched *vasa* signal. These four nuclei also exhibited PGC-like (diffuse) chromatin morphology by DAPI staining, as previously described. We conclude that the EICs are enriched for mRNAs of *vasa*, a second conserved PGC marker.

Taken together, the cell behaviors and *in situ* hybridization results above revealing enrichment of PGC marker mRNAs led us to conclude that the four EICs are the PGCs of *H. exemplaris*.

### *wiwi1* and *vasa* are present in developing oocytes and uniformly localized in early embryos

Since in other systems conserved markers of the PGC fate are often supplied maternally and then segregated to the PGC lineage, we performed FISH to examine the dynamics of *wiwi1* and *vasa* mRNAs in earlier developmental stages in *H. exemplaris* (Leclère et al., 2012; Mahowald, 1968; Marlow, 2010; Seydoux and Fire, 1994). Examination of developing oocytes within gravid adults required development of a FISH technique for adults. This proved technically challenging, since staining of adults requires bisection of the cuticle, as previously reported for immunofluorescence staining at this stage (Smith et al., 2017). Additionally, we found that oocytes would only stain when sliced open, as well, suggesting the presence of a permeability barrier preventing *in situ* reagents from reaching mRNAs inside oocytes (Supp. Fig. 2 C and Supp. Fig. 3 C). We observed the presence of *wiwi1* and *vasa* mRNAs in oocytes, suggesting that *wiwi1* and *vasa* mRNA are maternally supplied to embryos (Supp. Fig. 2 A and B and Supp. Fig. 3 A and B). We did not observe *wiwi1* and *vasa* mRNAs in surrounding, non-gonadal tissues in gravid adults.

We also performed FISH in one-cell and two-cell stage embryos. Similar to oocytes, *wiwi1* and *vasa* were present in one-cell and two-cell stage embryos (Fig. 4 A-A”, C-C” and Fig. 5 A-A”, CC”). Importantly, we did not see any polarized distribution, such as localization on one side of the embryo or in distinct regions of the embryo, which could otherwise reflect one means of restricting PGC markers to a subset of cells through development. Embryos between the two-cell stage and 12 hpl (cell internalization stage) could not maintain integrity through the *in situ* protocol, which requires manual removal of the eggshell. However, we obtained a small number of embryos at the four-cell and eight-cell stages that maintained partial integrity through the *in situ* protocol. Upon staining for *wiwi1* and *vasa* mRNAs by FISH, both *wiwi1* and *vasa* mRNAs were uniformly present in these embryos with no polarized distribution in one or more cells (Supp. Fig. 4). This suggests that at least through the two-cell stage (and possibly through the eight-cell stage), *wiwi1* and *vasa* mRNAs are uniformly present in all cells in *H. exemplaris* embryos.

**Figure 4.**
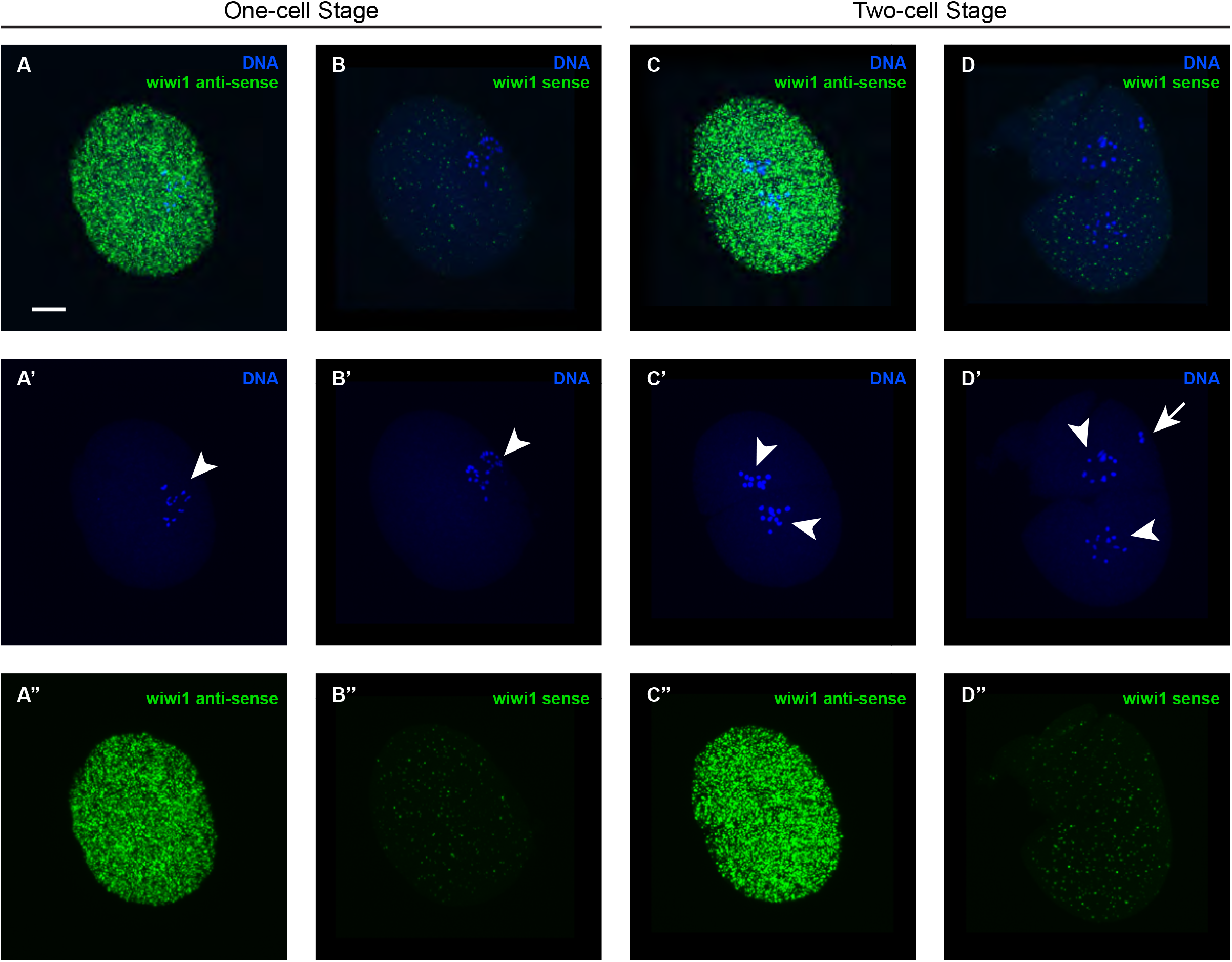
*wiwi1* mRNA is uniformly distributed in early embryos. (A-B) One-cell stage embryos. (C-D) Two-cell stage embryos. (A) Presence of *wiwi1* mRNAs by FISH in one-cell stage embryos. (C) Presence of *wiwi1* mRNAs by FISH in two-cell stage embryos. (n = 2 experiments of combined one and two cell stages, 8 and 7 embryos, respectively) Since, to our knowledge, this was the first published use of this technique in early stage embryos and since embryos at this stage are so yolk-dense, we included control embryos stained for the sense probe to the *wiwi1* mRNA, which showed little to no nonspecific signal. (B) Staining with *wiwi1* sense negative control probes shown for one-cell stage embryos. (D) Staining with *wiwi1* sense negative control probes shown for two-cell stage embryos. (n = 1 experiment of combined one and two cell stages, 6 embryos) Chromatin stained with DAPI is shown in blue, and *wiwi1* mRNA staining is shown in green. Arrowheads indicate nuclei. Arrow indicates polar body, which is only sometimes maintained through the protocol. (scale bar = 10 μm)

**Figure 5.**
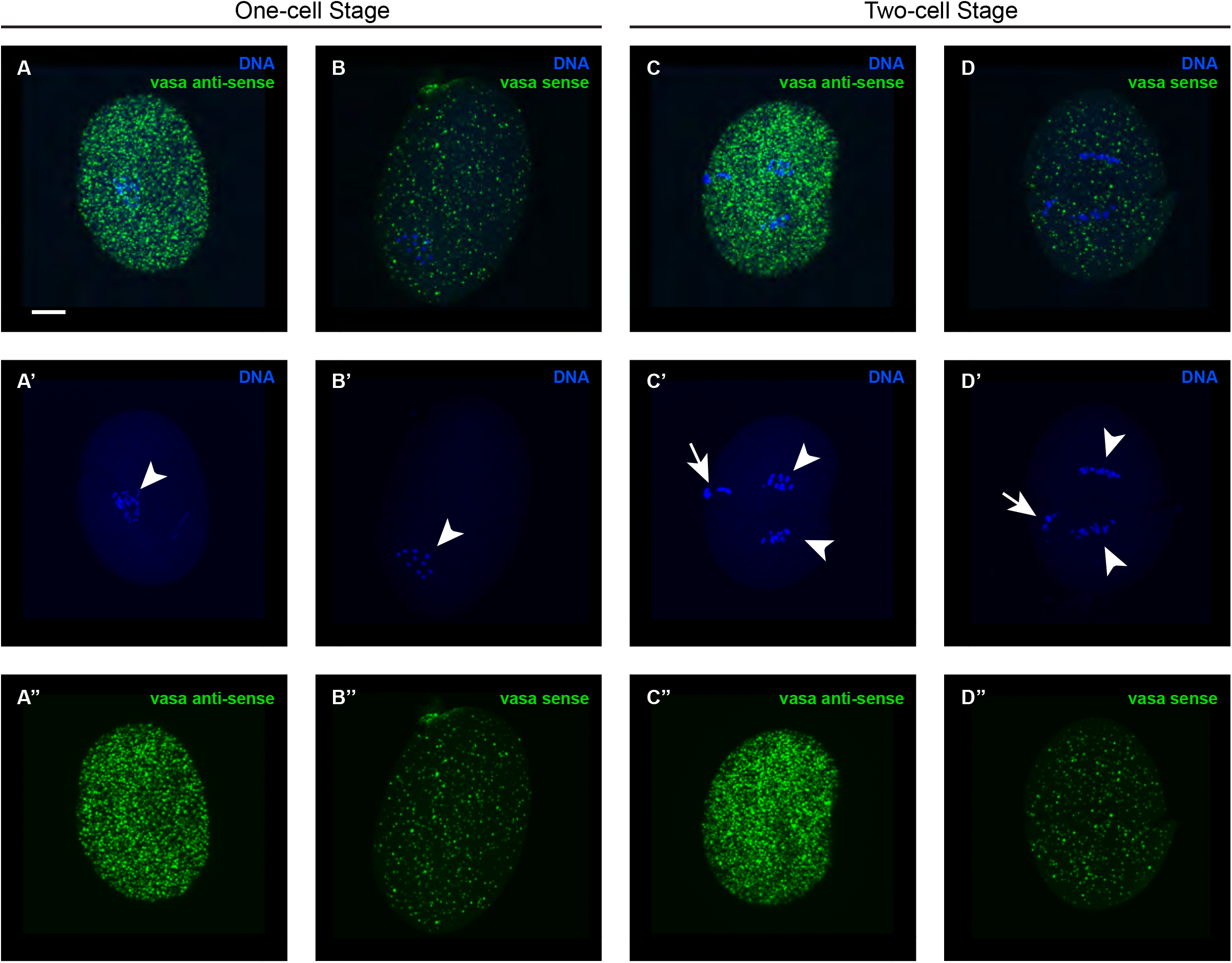
*vasa* mRNA is uniformly distributed in early embryos. (A-B) One-cell stage embryos. (C-D) Two-cell stage embryos. (A) Presence of *vasa* mRNAs by FISH in one-cell stage embryos. (C) Presence of *vasa* mRNAs by FISH in two-cell stage embryos. (n = 1 experiments of combined one and two cell stages, 6 embryos) Since to our knowledge, this was the first published use of this technique in early stage embryos and since embryos at this stage are so yolk-dense, we included control embryos stained for the sense probe to the *vasa* mRNA, which showed little to no nonspecific signal. (B) Staining with *vasa* sense negative control probes shown for one-cell stage embryos. (D) Staining with *vasa* sense negative control probes shown for two-cell stage embryos. (n = 1 experiment of combined one and two cell stages, 6 embryos) Chromatin stained with DAPI is shown in blue, and *vasa* mRNA staining is shown in green. Arrowheads indicate nuclei. Arrows indicate polar bodies, which are only sometimes maintained through the protocol. (scale bar = 10 μm)

### *wiwi1* and *vasa* exhibit dynamic enrichment through development

Since we did not see polarized distribution of *wiwi1* and *vasa* mRNAs in early embryonic stages, we sought to explain the enrichment of *wiwi1* and *vasa* mRNAs in the EICs. We examined the pattern of *wiwi1* and *vasa* mRNAs in the earliest time point after the 2-cell stage that could withstand the FISH protocol in our hands, 12 hpl. By this stage, the four EICs had internalized, and the embryo was still undergoing internalization of additional cells and had a developing epithelium evident surrounding the many internalizing cells (Fig. 1 A’) (Gabriel et al., 2007).

*wiwi1* mRNAs were present uniformly in all cells in embryos at 12 hpl (Fig. 6 A). This was also true at 13 hpl and at 14 hpl (Fig. 6 B and C). At 15 hpl there appeared to be a reduction in signal throughout the embryo (15 hpl and 12 hpl samples were stained in the same tube, to make approximate comparison of signal intensities possible) (Fig. 6 D). Signal remained low throughout the embryo at 17 hpl (13 hpl and 17hpl embryos were stained in the same tube), except in a small number of large cells at the posterior that showed an enrichment for *wiwi1* (Fig. 6 E). Similarly, at 18hpl, a subset of large cells at the posterior end exhibited enrichment for *wiwi1*, as expected from earlier results (Fig. 2 D), compared with the remainder of the embryo (14 hpl and 18 hpl embryos were stained in the same tube) (Fig. 6 F).

**Figure 6.**
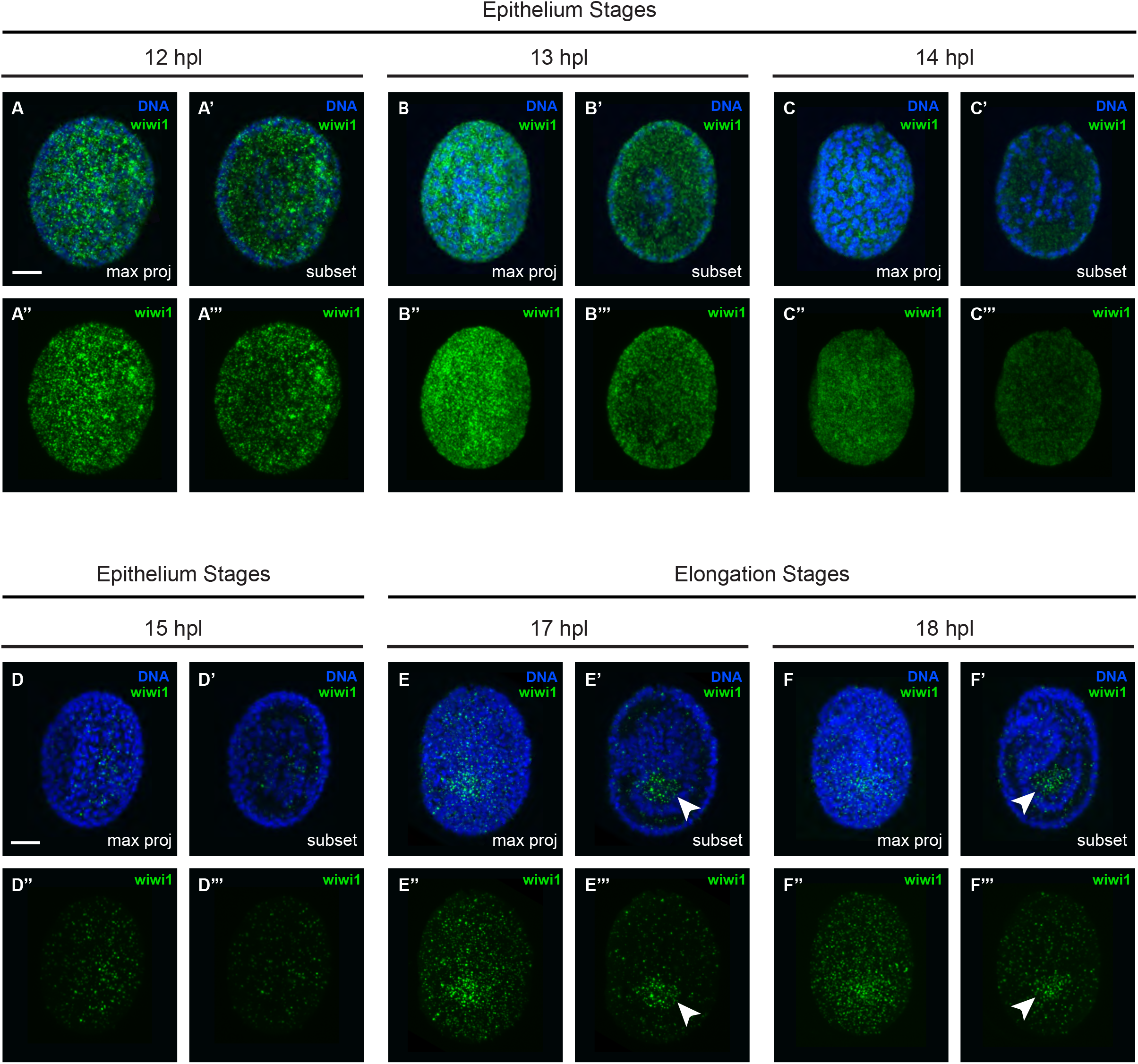
*wiwi1* mRNA exhibits dynamic enrichment through development. (A-F) Enrichment of *wiwi1* mRNAs through several stages of development. (A) 12 hpl (n = 2 experiments, 7 and 7 embryos, respectively). (B) 13 hpl (n = 2 experiments, 5 and 8 embryos, respectively). (C) 14 hpl (n = 2 experiments, 7 and 10 embryos, respectively). (D) 15 hpl (n = 2 experiments, 9 and 8 embryos, respectively). (E) 17 hpl (n = 2 experiments, 9 and 9 embryos, respectively). (F) 18 hpl (n = 2 experiments, 5 and 9 embryos, respectively). Maximum projections of total embryos shown on the left side for each time point and a projection of a subset of internal slices shown on the right side of each time point. Chromatin stained with DAPI is shown in blue, and *wiwi1* mRNA staining is shown in green. Arrowheads indicate the region enriched for *wiwi1* mRNA. (scale bars = 10 μm)

Conversely, *vasa* mRNAs exhibited enrichment in a small subset of cells starting at 12 hpl, although *vasa* was also present in the remainder of the embryo at lower levels at this stage (Fig. 7 A). This presence of *vasa* in all cells and enrichment in a subset of cells was also seen at 13 hpl and at 14 hpl (Fig. 7 B and C). At 15 hpl there appeared to be a reduction in signal throughout the embryo, except for a subset of cells that maintained a certain level of *vasa* mRNAs (15 hpl and 12 hpl samples were stained in the same tube) (Fig.7 D). Similarly to *wiwi1, vasa* signal remained low throughout the embryo except for a subset of large cells at 17 hpl (13 hpl and 17hpl embryos were stained in the same tube) (Fig. 7 E). This pattern was maintained at 18hpl, as expected from earlier results (Fig. 3 D, 14 hpl and 18 hpl embryos were stained in the same tube, Fig. 7 F).

**Figure 7.**
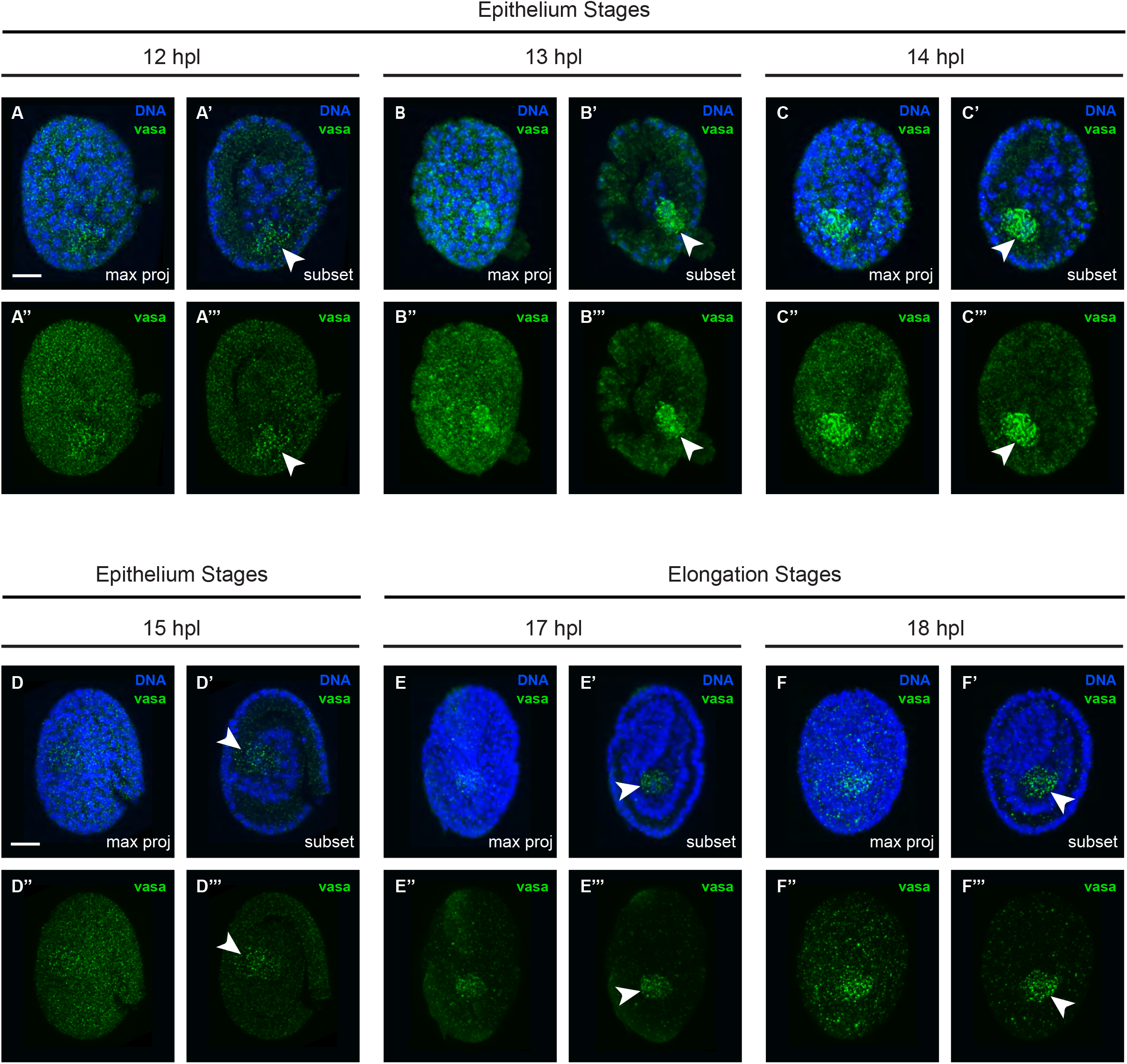
*vasa* mRNA exhibits dynamic enrichment through development. (A-F) Enrichment of *vasa* mRNAs through several stages of development. (A) 12 hpl (n = 2 experiments, 6 and 8 embryos, respectively). (B) 13 hpl (n = 2 experiments, 7 and 7 embryos, respectively). (C) 14 hpl (n = 2 experiments, 6 and 9 embryos, respectively). (D) 15 hpl (n = 2 experiments, 6 and 8 embryos, respectively). (E) 17 hpl (n = 2 experiments, 13 and 14 embryos, respectively). (F) 18 hpl (n = 2 experiments, 9 and 9 embryos, respectively). Maximum projections of total embryos shown on the left side for each time point and a projection of a subset of internal slices shown on the right side of each time point. Chromatin stained with DAPI is shown in blue, and *vasa* mRNA staining is shown in green. Arrowheads indicate the region enriched for *vasa* mRNA. (scale bars = 10 μm)

Considering the samples that were stained together, we could qualitatively extrapolate the dynamics of *wiwi1* and *vasa* mRNAs from 12 to 18 hpl. It appeared that *wiwi1* mRNAs were uniformly present throughout the embryo and decreased in level over time between 12 and 15 hpl, and then a small subset of large, undivided cells at the posterior end became enriched for *wiwi1* mRNAs by 17 hpl. This enrichment was maintained at 18 hpl. *vasa* mRNAs were present throughout the embryo at 12 hpl and were also enriched in a subset of cells. The *vasa* mRNAs decreased in levels over time between 12 and 15 hpl in all areas of the embryo, except for the subset enriched for *vasa*, and this subset of cells maintained its enrichment for *vasa* mRNAs through 17 and 18 hpl. Since samples earlier than 12 hpl did not remain intact through the FISH protocol, it remains an open question exactly when enrichment for *vasa* begins in a subset of cells. Since the position of the cells enriched for *wiwi1* and *vasa* at 17 and 18 hpl matches the position of the four large, undivided cells we defined above as the EICs, this led us to conclude that the EICs are also enriched for *wiwi1* and *vasa* mRNAs at these stages.

### The early cell lineage of *H. exemplaris* reveals the embryonic origin of the PGCs

To clarify the embryonic origin of the PGCs that we had identified in *H. exemplaris*, we constructed a cell lineage from the one-cell stage through 7 rounds of cleavages, and we noticed some discrepancies with the previously published lineage for this system (Gabriel et al., 2007). We filmed embryos from the one-cell stage through at least 20 hours of development by DIC microscopy, taking a z-stack every minute (stacks were 28-30 μm thick with 1 μm step size) (Fig. 8 A). We then collected nine of these films that were oriented such that the EICs could be followed throughout the film, and we tracked all cells and cleavage events manually using the FIJI plugin Mastodon (Pietzsch et al., 2020). We developed a program in MatLab to plot the mean (black horizontal lines) and standard deviation from the mean (blue error bars) times for each cell division, using the first cleavage event in each embryo as a starting time point (Fig. 8 B). While previously it was reported that three cells on the ventral side of the embryo undergo division arrest and internalize as the EICs, we found that four cells exhibited this behavior. Additionally, previously it was reported that the EICs arrested divisions after five rounds of division, but we found that the EICs arrest divisions after six rounds of division. Finally, while previously it was reported that the EICs arrested divisions before internalizing, we found some examples where one or both of the precursors to the EICs (anterior and posterior) internalized after only five rounds of division and then went through a sixth round of division after internalizing (Fig. 8 C). We observed the same nuclear migrations previously reported in the precursors to the four EICs, which suggest the divisions leading to the four EICs are asymmetric (Gabriel et al., 2007).

**Figure 8.**
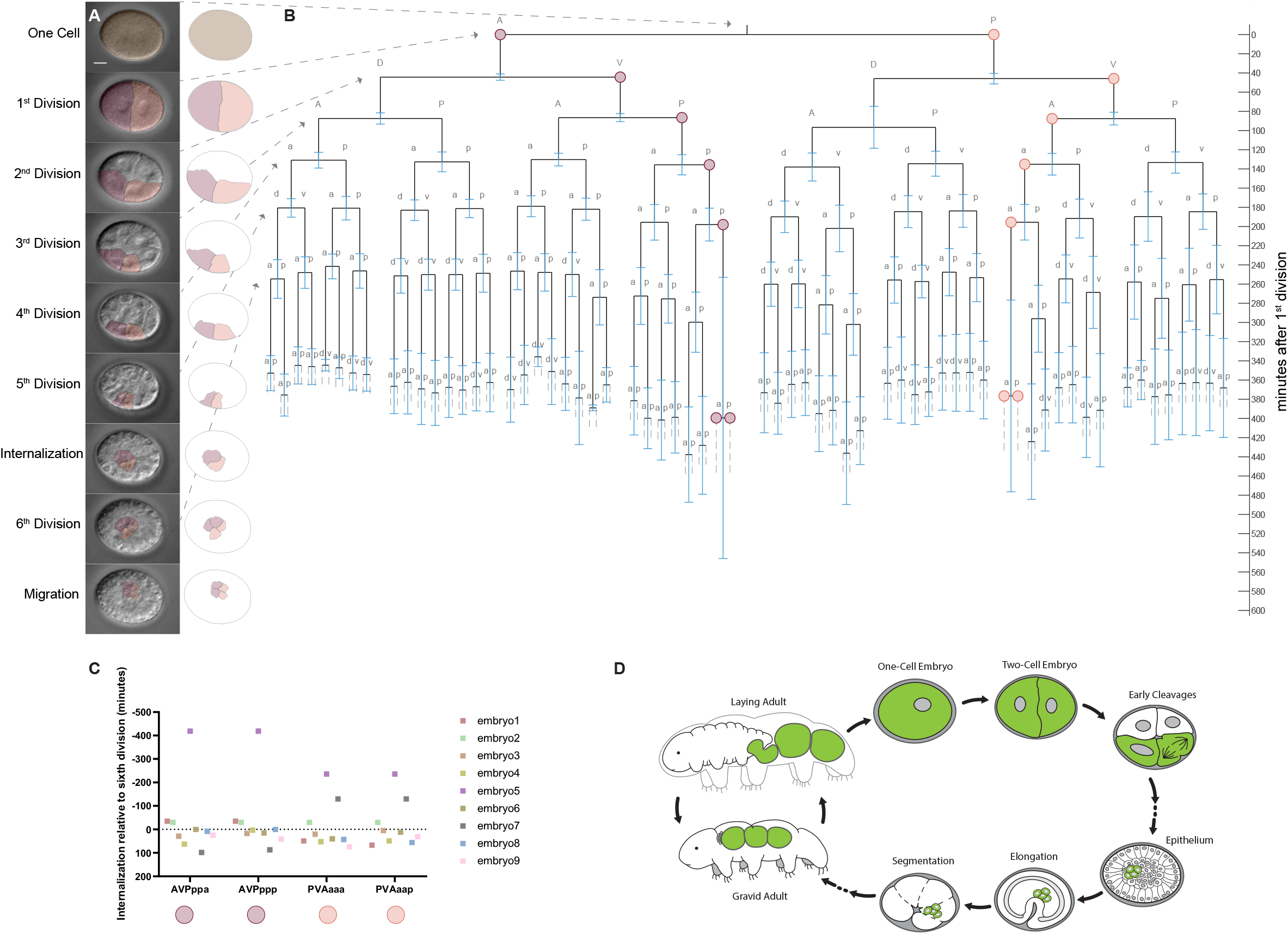
The early cell lineage of *H. exemplaris* reveals the embryonic origin of the PGCs. (A) Representative images of an early *H. exemplaris* embryo at nine timepoints: one-cell stage, first division, second division, third division, fourth division, fifth division, internalization of the EICs, sixth division, and migration of the four EICs (scale bar = 10 μm). Cells from the lineage leading to the four EICs are falsely colored with maroon and peach transparent overlays, for the anterior and posterior lineages, respectively. These overlays are shown in an outline of the embryo to the right of each image. (B) Lineage of the early *H. exemplaris* embryo, generated with division timing data from nine embryos, all followed from the one-cell stage through EIC migration (horizontal black bars are mean division times from 9 embryos, blue error bars indicate standard deviation from the mean). Division orientation is indicated by letters at each end of division mean time (a = anterior, p = posterior, d = dorsal, v = ventral). Axis along the right side of the lineage indicates the time of each division relative to the first division event, which was set at t = 0 for all nine embryos. Dotted arrows from A to B indicate the position of each representative image in A along the lineage. (C) Plot of internalization time (in minutes) for the four EICs relative to their sixth division, which is set at t = 0 (EICs are labeled along the horizontal axis by lineage name and with colored maroon and peach nodes, for the anterior and posterior lineages, respectively). Data is colored by embryo, and cells that internalized before undergoing the sixth division are shown at negative values above t = 0 on the inverted graph, while cells that internalized after undergoing the sixth division are shown at positive values below t = 0. Cells that were born into the middle of the embryo during the sixth division are shown at t = 0. (D) Drawing summarizing the traced embryonic origin of the PGC fate (in green) of *H. exemplaris*, through the stages in development observed: one-cell, two-cell, early cleavages, epithelium, elongation, segmentation, and gravid adult stages. Drawings modified from those by Heather Barber (Goldstein, 2022a), with permission.

Taken together with the data presented above, these results outline the embryonic origin of the PGCs in *H. exemplaris*. The PGCs arise on the ventral side of the embryo, from lineages that derive from each cell (anterior and posterior) of the two-cell stage embryo. After five rounds of division, the immediate precursors to the PGCs exhibit one of two behaviors: these cells either internalize first and then undergo a sixth division or undergo a sixth division first and then internalize. This stochasticity appears represented in and separable between the anterior and posterior lineages (Fig. 8 C). Based on these data and our identification of the PGCs above, we conclude that the four cells that are the earliest to internalize in the embryo are the four PGCs of *H. exemplaris*.

## Discussion

In this study, we revealed the fate of the four EICs of *H. exemplaris*. The EICs arise from lineages on both the anterior and posterior sides of the embryo, after six rounds of division, and are the first cells to internalize in the embryo. The EICs exhibit many conserved features of PGCs, including cell-cycle arrest, internalization from the embryo’s surface, migration to the location of the future gonad, and diffuse chromatin morphology. The EICs are also enriched for conserved markers of PGC fate: *wiwi1* and *vasa* mRNAs. These data collectively demonstrate that the four EICs are the PGCs.

Additionally, we characterized several aspects of PGC development in *H. exemplaris* by observing the dynamics of *wiwi1* and *vasa* mRNAs through different embryonic stages. By looking at the distribution of these mRNAs in one-cell and two-cell stage embryos, we found that cellspecific enrichment does not begin at these early stages, as mRNAs for both PGC markers were uniformly distributed, with no apparent localization. We found that the mRNAs of *wiwi1* become enriched in the PGC lineage between 15 and 17 hpl, during embryonic elongation, with a global reduction in *wiwi1* prior to this. The mRNAs of *vasa* become locally enriched sooner in the PGC lineage between the two-cell stage and 12 hpl (the epithelium stage) with a global reduction in *vasa* between 14 and 15 hpl. Because embryos between the two-cell stage and 12 hpl do not maintain integrity through the *in situ* protocol, we were unable to deduce exactly when *vasa* enrichment begins in development. The few embryos at the four-cell and eight-cell stages that remained intact through the *in situ* protocol revealed that it is likely *wiwi1* and *vasa* are also not enriched in the PGC lineage at these stages. Therefore, it is likely that *vasa* mRNAs begin to be enriched in the PGC lineage sometime between the eight-cell stage and 12 hpl. Development of a gentler *in situ* protocol (with a gentler eggshell removal method) will be necessary to stain embryos between these stages and deduce when exactly *vasa* mRNA enrichment in the PGC lineage begins. It is possible that *wiwi1* mRNAs are transiently enriched in a subset of cells within this window of time during which *in situs* are not possible and then are more uniformly distributed across the embryo again by 12 hpl. If such dynamics exist, we would have missed them due to the limited stages at which *in situs* are feasible. We were able to detect *wiwi1* and *vasa* mRNAs in developing oocytes in gravid adult tardigrades upon application of the embryo *in situ* protocol to adults. This result indicates that PGC marker mRNAs are maternally provided to embryos prior to egg-laying, during oogenesis. We never observed staining for PGC marker mRNAs in germ stem cell niche cells in the germarium, evident as an arc of small cells sitting at the anterior end of the gonad (Jezierska et al., 2021), even when the germ stem cell niche was located directly adjacent to a cut oocyte that showed staining. This could be because these mRNAs are not present in the germ stem cell niche or because the germ stem cell niche is surrounded by an additional barrier that must be further permeabilized to the *in situ* reagents.

The temporal dynamics of these PGC marker mRNAs revealed that they are primarily enriched in the PGC lineage. This did not necessarily need to be the case, as in some systems, embryonic enrichment is seen first for the proteins of PGC markers and either second or not at all for their mRNAs, as in the urchin *Lytechinus variegatus* and the nematode *C. elegans* (Fresques et al., 2016; Voronina et al., 2011). It is possible that we have missed an earlier enrichment of these PGC markers to the PGC lineage at the protein level, through translational or post-translational regulation. However, sometimes mRNA enrichment precedes protein enrichment in PGCs, as in the zebrafish *Danio rerio* (Knaut et al., 2000). Since we did see enrichment of the mRNAs for these PGC markers, we can at least conclude that these markers are enriched at the mRNA level to the PGC lineage by 12 hpl (for *vasa* mRNA) and 17 hpl (for *wiwi1* mRNA). Our attempts to use cross-reactive antibodies to Vasa protein were not successful (see Methods). Additionally, many studies have made clear that these PGC marker mRNAs are present in certain somatic and multipotent stem cells (Juliano et al., 2010). However, because the cells enriched for these mRNAs in *H. exemplaris* were the same cells that exhibited PGC-like behaviors, this led us to conclude that in the stages observed, *vasa* and *piwi* mRNAs mark the PGCs.

We were unable to identify an *H. exemplaris* homolog of one of the highly conserved PGC markers: *nanos*. We also failed to identify a *nanos* homolog in another tardigrade species, *R. varieornatus*. Previously, *nanos* was found to have a slightly elevated rate of sequence evolution among species of *Drosophila*, compared with other germ cell markers (Whittle and Extavour, 2019), and thus the failure to identify a homolog of *nanos* among tardigrade species could be because of low sequence conservation, because a tardigrade homolog of *nanos* does not exist, or because the published sequence databases are incomplete and each failed to capture a *nanos* homolog. If there is not a tardigrade homolog of *nanos*, then it is possible another gene is compensating for the lack of *nanos*.

The mRNA enrichment dynamics observed lead to the question: how do PGC markers become enriched in a subset of all cells in *H. exemplaris?* We considered several possibilities. First, mRNAs could be degraded in all cells, with mRNA expression occurring afterwards in a small subset of cells, as with *vasa* mRNA in *D. melanogaster* embryos (Hay et al., 1988). Second, mRNAs could be degraded in all except for a small subset of cells, which retain mRNAs, as with *vasa* mRNA in *Oryzias latipes* (medaka) embryos (Herpin et al., 2007; Shinomiya et al., 2000). Third, mRNAs could be shuttled from all cells to a small subset through membrane nanotubes, extracellular vesicles, or other cytoplasmic connections between cells (Haimovich et al., 2017; Valadi et al., 2007). Fourth, some combination of degradation, increased expression, maintenance, and shuttling of mRNAs is possible. Distinguishing between these possibilities will require the development of techniques not yet available in this system, particularly means of cell-specific manipulation and of labeling cellular components for live fluorescent imaging. Work toward the latter has begun with the use of different live fluorescent dyes to temporarily label various cellular components, such as mitochondria, lysosomes, and membrane (McGreevy et al., 2018). Transgene and CRISPR techniques were recently developed for somatic cells of adult tardigrades, but they have not yet been demonstrated to work in germline cells (Kumagai et al., 2022; Tanaka et al., 2022). There has yet to be a technique established for labeling and live imaging of specific mRNA and/or proteins or for live reporting of mRNA transcription and/or translation in germline or embryos of this system.

This work lays the foundation for future studies to determine how PGCs are specified in *H. exemplaris*. Known mechanisms of PGC specification in metazoans are often classified in either of two modes: inheritance and induction (Extavour and Akam, 2003; Whittle and Extavour, 2017). The inheritance mode of PGC fate involves sequestration of maternally supplied determinants and/or protectants to germ granules, which are segregated to invariant cell lineage(s) by cytoplasmic partitioning during successive rounds of cell divisions, as seen in the partitioning of p granules in *C. elegans* embryos (Strome and Wood, 1983). The inductive mode uses many of the same mRNAs and proteins, but instead, these are expressed zygotically, with expression and maintenance restricted to certain cells that have received a cue to assume the PGC fate by signaling from other cells in the embryo, as with PGC specification in *Gryllus bimaculatus* (Ewen-Campen et al., 2013). The two modes of PGC specification are seen throughout the metazoan tree, and the inductive mode is hypothesized to be the ancestral mode of PGC specification for the metazoa (Extavour and Akam, 2003; Juliano et al., 2010; Whittle and Extavour, 2017).

Both modes of specification are found in the protostome superclade ecdysozoa, which includes the phyla nematoda, arthropoda, and tardigrada. PGCs are specified by the inheritance mode in the nematode *Caenorhabditis elegans* (roundworm) and the arthropod *Drosophila melanogaster* (fruit fly) and by the inductive mode in the arthropod *G. bimaculatus* (crickets) (Ewen-Campen et al., 2013; Extavour and Akam, 2003; Strome, 2005; Wang and Seydoux, 2013; Whittle and Extavour, 2017; Williamson and Lehmann, 1996). To our knowledge, there has yet to be a nematode species whose PGC specification mode has been identified as the inductive mode (Whittle and Extavour, 2017). A recent study found that features of both modes may be present in the formation of PGCs in *D. melanogaster* (Colonnetta, et al., 2022). Further knowledge of PGC development and specification in ecdysozoan phyla is needed to gain a clearer view of the evolution of PGC specification mechanisms within this superclade. From our work here, we cannot deduce which mode is used in the tardigrade *H. exemplaris*. Expression patterns of *wiwi1* and *vasa* indicate there is no cytoplasmic partitioning of these mRNAs at early stages. The dynamics of these mRNAs are similar to those of *vasa* and *piwi* mRNAs in *Patiria miniata* embryos, which undergo inductive germ cell specification (Fresques et al., 2014; Fresques and Wessel, 2018). Additionally, previous electron microscopy of early *H. exemplaris* embryos failed to find evidence of a germ plasm, as is seen in systems that use the inheritance mode (Gabriel, 2007). Therefore, these data appear more consistent with the hypothesis that the inductive mode is used in *H. exemplaris* PGC specification. Future studies are needed to clarify the mode of PGC specification in this system. It may be that *H. exemplaris* PGC specification occurs using a mix of features from the two modes.

Previous studies have hypothesized that PGCs are specified by inductive modes in various tardigrade species, based on observations of embryos by light microscopy (Extavour and Akam, 2003; Marcus, 1929; von Erlanger, 1895; von Wenck, 1914). These studies labeled germ cells (‘urkeimzellen’ or ‘keimbläschen’ in German) based on their morphology and positioning in the embryo. Our observations build on these previous studies by adding molecular evidence of PGC fate to precisely label the PGCs and manual tracing of the embryonic lineage that leads to the PGCs. One study found that another tardigrade species *Thulinia stephaniae* is able to compensate for the loss of half of the embryo by laser ablation at the two-cell or four-cell stage and develop normally (albeit smaller), including normal germ cell specification (Hejnol and Schnabel, 2005). Upon ablation, the correct number of PGC-like cells (identified by cellular size and behavior) was observed in the expected position. This result suggests that PGC specification is regulative and does not depend upon specific cytoplasmic partitioning of germplasm. However, other studies have revealed organisms whose germlines normally are specified with cytoplasmic partitioning of germ granules, but that also have the potential to regulate the loss of the typical germline and induce a new germline later in development, such as sea urchins (Yajima and Wessel, 2011). Therefore, we believe the ablation data presented in this study reveals the regulative potential of PGCs in *T. stephaniae* embryos but may not reveal what happens normally in development. It remains an open question what modes of PGC specification exist in the tardigrade phylum.

Through lineaging, we discovered that the PGCs are set aside after six rounds of division, at which point they undergo cell cycle arrest. Although we did not follow them further into development, we anticipate that this cell cycle arrest is temporary and that the PGCs resume divisions later to establish the germline, as happens in other systems (Su et al., 1998).

We reported several differences between the previously published lineage and the lineage we observed in *H. exemplaris* embryos (Gabriel et al., 2007). We were unable to determine the reason for these differences, but several possibilities exist. First, the animal population used to establish the previous cell lineage could be a slightly different strain and/or species from the one we studied. Second, the species could be the same and exhibit different lineages either due to differences in rearing conditions or due to rapid evolution in the time between curation of the lineage previously published in 2007 and our lineage. Third, the previously-published lineage could have some errors due to missed cell cycles and/or cell internalizations. We were unable to recover the videos used to build the previously-published lineage and hence, cannot identify the cause of the differences we see. Regardless, the lineage presented in this paper reflects the lineage of *H. exemplaris* in our hands and reveals when the PGCs arise in their development.

## Conclusions

This work is the first molecular characterization of an early cell lineage in *H. exemplaris* and in the tardigrade phylum more broadly. By identifying the origin of the PGC lineage in *H. exemplaris* embryos, this work lays the foundation upon which further studies may explore the mechanism of PGC specification in this animal and more broadly, in this phylum. Comparing this to known mechanisms of PGC specification in related and distant phyla will aid in building our understanding of how mechanisms of PGC specification have evolved.

## Materials and Methods

### Maintaining Cultures of *Hypsibius exemplaris*

Cultures of *Hypsibius exemplaris* were maintained as previously described (Mcnuff, 2018). Briefly, animals were kept in 3.5 cm plastic petri dishes (VWR product # 102097-174) with approximately 2.5 mL Deer Park brand spring water and approximately 0.5 mL Chloroccocum sp. algae. Cultures were split approximately every week from month-old cultures and kept in a plastic box with wet paper towels lining the bottom to maintain humidity and prevent evaporation.

### DIC Imaging of Development

Embryos were imaged by DIC light microscopy as previously described (Heikes and Goldstein, 2018b). Briefly, embryos were mounted on slides with 15 μL Deer Park spring water and 28.41 μm glass beads (Whitehouse Scientific product no. MS0028) and covered with No. 1.5 coverslips, the edges of which were sealed with VALAP. Each slide was then imaged on a Nikon Eclipse e800 microscope with a 100x oil objective and a C2400-07 Hamamatsu Newvicon camera or a pco.Panda 26 camera. Images were acquired using Metamorph (Molecular Devices). Stacks were taken every minute for at least 24 hours (or over several acquisitions for a total of 24 hours) with a z-step size of 1 μm for a total of 28-30 μm per embryo. Some drift in z was inevitable, so acquisitions were often paused and resumed after some adjustment in z to ensure minimal loss of information due to the stage drifting away from the objective. Metamorph saved raw images as .tiff files with an .nd file to instruct software on how to open the images. To speed up analysis, images were opened in FIJI (Schindelin et al., 2012), compressed to 8-bit, and then saved as a copy as multidimensional .tiff files. These files were used for lineaging. Raw images were opened and still frames were isolated and saved as .eps files for the examples shown in Figures 1 and 8. These .eps files were further annotated in Adobe Illustrator to outline cells of interest. We note that we observed natural heterogeneity in embryo sizes, though they were all approximately close to the size previously reported (Gabriel et al., 2007). Experimental causes of size variability include variable degree of compression between slide and coverslip, even with the glass beads used to prop the coverslip. Additionally, embryo sizes may appear varied in images, depending on the z position of the representative images used in figures; positions closer to the center plane of the embryo will appear larger than positions closer to the objective.

### Manual Lineaging of Early Development with Mastodon

Lineaging was performed in the FIJI (Schindelin et al., 2012) plugin Mastodon (Pietzsch et al., 2020). Briefly, a .xml file was saved for each embryo .tiff film using BigDataViewer, with the command Plugins>Multiview Reconstruction>Batch Processing>Define Multiview Dataset (Pietzsch et al., 2015). The dataset type selected was “Zeiss Lightsheet Z.1 Dataset (LOCI Bioinformats)” and the file name was given a .xml suffix. The image location was entered as the .tiff file for the dataset, and all default settings were used before saving the .xml file. This .xml file was then used to open the dataset in Mastodon and save it as a mastodon file. Lineaging was performed manually by following cell nuclei (the location in cells where the DIC shows yolk exclusion). The approximate center of each nucleus was labeled with a point, and when a cell divided, the division was marked by linking the center point of the mother cell’s nucleus in one time point to the center points of the two daughter cells’ nuclei in the following time point. The completion of a cell division was defined as apparent completion of cytokinesis that could be seen to the best of our ability by a clear, thin boundary of cell membrane (and potentially yolk and cytoplasm) between recently-divided nuclei. Daughter cells after each division were named by division orientation relative to the axes of the embryo. For cells that divided in an oblique orientation, daughter cells were named by their relative position in at least two of the three body axes. Cells were tracked in embryos for as long as their nuclei were apparent. When a cell moved to a location too far from the coverslip for its nucleus to be seen clearly, then it was no longer followed (hence, some cells were followed in some but not all films). Cells were tracked from the beginning of each .tiff film through seven rounds of division, or until nuclei were no longer clearly visible, except for the four putative PGCs; these lineages were tracked for six rounds of division until the four cells were born and internalized and began to migrate. Each division timepoint was saved digitally in the Mastodon file and also manually recorded on paper in a hand-written version of the lineage. These data were then used to plot the lineage in MATLAB (MathWorks). Lineaging was performed on .tiff films from 9 embryos.

### Plotting Manual Lineaging in MATLAB

Lineaging results from the 9 embryos were compared and given common names for each cell, by most common division orientation. After this, a mean normalized division time was calculated for each cell across the 9 embryos. Each cell had at least 5 replicates represented amongst the dataset through 6 rounds of division. The 7th round of division included at least 3 replicates for each cell, except for AVAppp and AVAppa, which only divided in a clear enough view in 2 of the embryos analyzed. This is due to these cells often being positioned too far from the coverslip. Standard deviation from the mean was also calculated for each cell. These values were all calculated and saved in an Excel spreadsheet. A MATLAB (MathWorks) program was developed for plotting the lineage data. The program performs the following actions: plot the mean lineage of embryos using the mean of each division event across embryo samples, overlay standard deviation from the mean at each division, label division orientations, and highlight key features of the lineage (in this case, highlighting the nodes of the lineage leading to the four EICs). This program is not species-specific and only requires that cells be labeled by the orientation of divisions leading to their existence (dorsal-ventral, anterior-posterior, and left-right; e.g. PV is the ventral cell arising from the division of the P (posterior) cell in the dorsal-ventral axis of the embryo). We developed this in MATLAB version R2019a and called it ELM (Embryonic Lineage in Matlab), after the tree. This program is freely available via Github at: https://github.com/kiraheikes/ELM. The program read the data from the Excel spreadsheet that we curated and plotted each average division time as a horizontal line and each new cell arising from each division as a vertical line. Beyond the seventh round of division (sixth for the four EICs), dashed lines were plotted to indicate that these cells continue to exist, but the remainder of these cells’ lineages were not tracked. The nodes of the cells leading to the four EICs were marked with colored nodes (maroon and peach for the anterior and posterior lineages, respectively). The four EICs were also marked with these colored nodes at the end of each to indicate that these cells arrested divisions at this time. Standard deviation from the mean was plotted with blue bars at each division time. Internalization times for the four EICs were plotted separately from the cell lineage plot, as these varied quite a bit relative to the sixth division time for each cell amongst the 9 embryos. This plot was generated in Prism. A dotted horizontal line at t=0 was included to indicate the time of the sixth division for each of the four cells, and each dot plotted represents the time of internalization relative to division time for each of the four cells from each embryo.

### Gene Identification and Phylogenetic Analyses

Genes were identified by tBLASTn (Altschul et al., 1990; Gerts et al., 2006; Sayers et al., 2022) using protein sequences for known homologs from *Drosophila melanogaster* found in the UCSC genome browser (Kent et al., 2002) against the *Hypsibius exemplaris* transcriptome published in (Levin et al., 2016). A reciprocal BLASTp was performed with the top hits against the *D. melanogaster* transcriptome (Sayers et al., 2022). Top hits that successfully returned the original query by reciprocal BLAST were further analyzed using the NCBI conserved domain analysis tool (Lu et al., 2020; Marchler-Bauer et al., 2017, 2015, 2011; Marchler-Bauer and Bryant, 2004) and SMART domain analysis tool in normal mode (Letunic et al., 2021; Letunic and Bork, 2018) to confirm that the protein structure matched that of the query. Maximum likelihood phylogenetic reconstructions were produced comparing the top hits for each gene to known homologs. First, homologous protein sequences were aligned by the MUSCLE algorithm using MEGA-X software (Kumar et al., 2018). These alignments were then trimmed with the GBLOCKS server developed by the Castrasena lab using the least stringent selection (Castresana, 2000; Dereeper et al., 2008). These trimmed alignments were then used to construct a Maximum Likelihood tree in MEGA-X using the Maximum Likelihood method and Whelan and Goldman model with 500 bootstraps (Felsenstein, 1985; Kumar et al., 2018; Whelan and Goldman, 2001). The percentage of replicate trees in which the associated taxa clustered together in the bootstrap test (500 replicates) are shown next to the branches. Initial tree(s) for the heuristic search were obtained automatically by applying Neighbor-Joining and BioNJ algorithms to a matrix of pairwise distances estimated using a JTT model, and then selecting the topology with superior log likelihood value. Final trees were exported as .eps files and labels were added using Adobe Illustrator. Table 1 lists the accession numbers for homologous protein sequences used in the alignments for each gene, similar to those used in (Ewen-Campen et al., 2013) to identify the sequences for *Gryllus bimaculatus* Piwi and Vasa. Protein domains were identified with NCBI’s conserved domain search. Multiple Sequence Alignments of amino acid sequences of domains were colored in Jalview by Percentage Identity (<40%, >40%, >60%, and >80%, from white thru darker shades of gray) (Waterhouse et al., 2009). Color was adjusted in Adobe Illustrator.

### Cloning

Genes were cloned as previously described (Smith et al., 2016). Primers were designed using NCBI PrimerBLAST (Ye et al., 2012) to amplify each gene in a series of two nested PCRs and isolate a product that was 700-800 bp in length, which is an optimum length for *in situ* hybridization probes. First rounds of PCR reactions were performed with GoTaq polymerase and the primers using *H. exemplaris* total embryo cDNA, generated from hundreds of embryos and a few dozen adults spiked in, as a template. Second rounds of nested PCR reactions were performed with GoTaq polymerase, using the products of the first reaction as a template. Products were run on 2% agarose gels to confirm amplification of sequences of the expected lengths. Amplified products were then cloned into pCR 4-TOPO TA vectors (Invitrogen, with the exception of *wiwi1*, which was cloned using blunt-end cloning into a pCR 4Blunt-TOPO vector, as insertion into the pCR 4-TOPO TA vector proved tricky). NEB DH5-alpha cells were mixed with 2 μL of pCR TOPO vector reaction and were transformed and grown according to the NEB protocol (https://www.neb.com/protocols/0001/01/01/high-efficiency-transformation-protocol-c2987).

Cells were plated on LB-Ampicillin selection plates and grown overnight. Colonies were picked and streaked and grown on LB-Ampicillin selection plates and also used as templates for colony PCR to confirm vector insertions at the expected sizes, using optimized versions of the M13 primer sequences. Cells from colonies with proper vector insertion sizes were then grown in 5-mL falcon tube cultures with LB-Ampicillin, shaking overnight to amplify cells containing the vectors with isolated products. Cultures were spun-down and miniprepped and submitted for sequencing by Genewiz. Sequence results were analyzed and aligned to the expected sequence for each gene. Colonies were selected for probe synthesis based on sequencing results with the least number of changes from the published, expected sequence. Accession numbers for the gene sequences used are listed in Table 1. Primers are listed in Table 2.

### Probe Synthesis

RNA *in situ* hybridization probes were synthesized as previously described (Smith, 2018; Smith et al., 2016), using the T7/T3 Riboprobe Combination System kit (Promega) and the Digoxigenin labeling kit (Roche). Both anti-sense and sense probes were made for each gene, using T7 or T3 for each, depending on the orientation of each gene insertion in the TOPO vector (identified by sequencing results). Probes were cleaned using an RNEasy MinElute Cleanup Kit (Qiagen) and diluted in RNAse-free water and kept at −80C. Final probe concentration for *in situ* reactions was 0.5 μg/mL, as recommended in (Smith, 2018).

### Chromogenic *In Situ* Hybridization

Chromogenic *in situ* hybridization was performed as previously described (Smith, 2018). Briefly, embryos were collected in a dish with spring water at the 1-cell stage (typically still sharing the exuvia with the adult) and were left to age to an hour and fifteen minutes before the desired age. Then, embryos were cut from exuvia and transferred to a 1.5 mL tube to permeabilize in chitinase and chymotrypsin for an hour. Early-stage (one and two-cell) embryos were only permeabilized for 20-30 mins. The chitinase and chymotrypsin solution was washed out and embryos were then fixed in the 1.5 mL tube in fixative solution (as described) for 30 minutes, rocking gently. Fixative was washed out and embryos were then transferred to a mobicol column (as described) and dehydrated in a gradual methanol series from 25% to 100%. Embryos were stored at −20°C overnight to dehydrate. Embryos were then rehydrated in a gradual series from 100% methanol to 100% 0.5X PBTween. Embryos were transferred to a dish with 0.5X PBTween using a glass pasteur pipette. Embryo eggshells were cut using a syringe needle, and then embryos were recollected into new mobicol columns, one for each probe of interest. Embryos were washed with 1% triethanolamine and then acetylated with 0.2% acetic anhydride in 1% triethanolamine twice, for five minutes each. Acetic anhydride solution was washed out, and then embryos were prehybridized with 50% hybe solution (as described) and 50% 0.5X PBTween, rocking gently for 20 minutes. Embryos were then prehybridized with 100% hybe solution, rocking gently for 20 minutes. Then, embryos were prehybridized with 100% hybe solution heated to 60°C for 2 hours, kept in a heat block set to 60°C. Embryos were then incubated with 0.5 μg/mL probe in hybe solution overnight (at least 16 hours) in the heat block set to 60°C, covered in foil. Sense negative control probes were used for every time point and every gene examined at least once. Hybe solution was washed-out with plain hybe solution (as described) heated to 60°C, 5 times quickly and 5 times for 20 minutes each time. Embryos were then washed in an SSC solution series, as described. Embryos were washed quickly with 0.5X PBTween two times. Embryos were incubated in blocking solution (as described) for two hours at room temperature, rocking gently, covered in foil. Then, embryos were incubated with 1:1500 anti-DIG::POD antibody (Roche) in blocking solution overnight at 4°C, rocking gently, covered in foil. Antibody was washed out with MAB solution, five times quickly and five times for 10 minutes. Embryos were then washed with AP development solution (as described) three times quickly and one time for ten minutes. Embryos were kept covered in foil through these washes. Embryos were then transferred to a glass depression slide using a glass pasteur pipette and were collected under a dissection microscope and transferred to a well of a 96-well plate filled with 150 μL of BM Purple development solution (Roche). The 96-well plate was kept in the dark for embryo staining to develop. Once developed, the reaction was quenched by transferring embryos to a new well in the plate with 150 μL 0.5X PBTween. Plates were kept at 4°C in a box with wet kimwipes until ready to image.

### Widefield Imaging of Chromogenic *In Situs*

CISH-stained embryos were transferred with a glass pasteur pipette to a depression slide filled with 0.5X PBTween and imaged using a EOS Rebel T6 Canon camera mounted to a Zeiss Axiozoom V.16 microscope with a Plan-NeoFluar Z 1x/0.25 FWD 56mm objective lens. Images were taken remotely using Canon EOS 3 software on a laptop connected via usb to the camera. Anti-sense experimental and sense negative control samples were imaged back-to-back using the same settings.

### Fluorescent *In Situ* Hybridization

Fluorescent *in situ* hybridization was performed using the Tyramide Signal Amplification kit (Akoya Biosciences). A more detailed protocol can be found in the Supplement. All steps were similar to the CISH protocol described above and published (Smith, 2018), except for a few key differences. First, the post-hybridization washes were performed in a quicker manner. Specifically, embryos were washed ten times quickly with plain hybe solution, six times quickly with 2x SSC solution, two times quickly with 1x SSC solution, and two times quickly with 0.2x SSC solution, before continuing to the two quick washes with room temperature 0.5X PBTween. Second, the embryos were blocked with the blocking buffer that is included with the TSA Signal Amplification kit from Akoya Biosciences, again rocking for two hours at room temperature. Third, the DIG probes were detected with a different antibody: Anti-DIG-POD, which was diluted 1:500 in the same blocking buffer that is included with the TSA Signal Amplification kit from Akoya Biosciences. Embryos were incubated with this overnight, rocking gently at 4°C. Fourth, after washing-out the antibody, the embryos were washed for five minutes in 500 μL of the 1X amplification diluent included in the TSA Signal Amplification kit from Akoya Biosciences, rocking at room temperature. Fifth, instead of developing the *in situs*, the DIG-labeled probes were amplified with the Cy3 TSA reagent included with the TSA Signal Amplification kit from Akoya Biosciences, diluted 1:50 in the same 1X amplification diluent in a 200 μL solution. This was then washed-out with 0.5X PBTween five times quickly and five times for ten minutes, all heated to 60°C. Samples were kept in the mobicol columns at 4°C until ready to mount. Adults were stained using the same protocol. To prepare adults for staining, they were prepared as described for immunostaining (Smith et al., 2017), but with 0.5X PBTween instead of 0.5X PBTriton. Briefly, adults were collected while visibly gravid and transferred to seltzer water to stretch-out. They were then fixed in a 1.5 mL tube with 4% paraformaldehyde in 0.5X PBTween. Animals were stored in this solution at 4°C at least overnight until ready to stain. To permeabilize, animals were washed five times for five minutes with 0.5X PBTween and then transferred to a dish with 0.5X PBTween. Animals were bisected with a 25-gauge syringe needle. We found that oocytes would only stain when they were also cut during this bisection. Animals were then transferred into mobicol columns and taken through the same *in situ* protocol as the embryos. Sense negative control probes were used for every time point and every gene examined at least once.

### Immunofluorescence

Immunofluorescence was performed as previously described (Smith and Gabriel, 2018). We attempted to examine Vasa protein distribution using a cross-reactive antibody to Vasa (Sgv, *Schistocerca gregaria vasa*) that was kindly gifted to us by Dr. Gee-Way Lin and Dr. Chun-Che Chang and that has worked in diverse arthropods (personal communication, Formosa2 antibody was unavailable). Although we were able to repeat the use of this antibody to stain Vasa protein in PGCs in *D. melanogaster* embryos, we were unable to see any staining in *H. exemplaris* embryos with this antibody (data not shown). We also attempted to use another cross-reactive antibody to Vasa (Lasko and Ashburner, 1990) that was kindly gifted to us by Dr. Hong Han and Dr. Paul Lasko, but we were unable to repeat staining of Vasa in PGCs in *D. melanogaster* or to see any staining in *H. exemplaris* embryos (data not shown).

### Fluorescent Imaging

FISH-stained embryos and adults were mounted in 2 μL DAPI fluoromount-G (Southern Biotech) with 37.36 μm glass beads (Whitehouse scientific) on slides and covered with No. 1.5 coverslips. Slides were allowed to cure overnight in the dark at room temperature, and then coverslip edges were sealed with a fast-drying clear nail polish. Samples were imaged on a Zeiss 710 LSM with a Plan-Neofluor 100x/1.3 oil Iris objective (for embryos) or a Plan-Apochromat 63x/1.40 oil DIC M27 objective (for adults). DAPI signal was excited with a 405 nm laser and collected between 418 and 570 nm, and Cy3 signal was excited with a 560 nm laser and collected between 581 and 695 nm. Images were analyzed in Zen Black (Zeiss).

### Curating Videos in Imaris

Fluorescent videos were curated in Imaris (Oxford Instruments). Fluorescent data was displayed as a maximum intensity projection. Manually-cropped regions were made by creating a new Imaris surface and then drawing around the highly enriched region for each mRNA (*wiwi1* and *vasa*) slice-by-slice. The resulting stack of manually-drawn regions were stitched together by Imaris into a surface. This was used to mask the full maximum intensity projection data into a maximum intensity projection of only the cropped region. The masked DAPI data was then segmented by creating a new Imaris surface and assigning the following thresholding for the *wiwi1* and *vasa* videos, respectively. For the *wiwi1* video, surfaces were seeded with a grain size of 0.2 μm and largest sphere diameter of 2.0 μm. These were thresholded to an intensity value of 22.9817 a.u., and Imaris was instructed to split connected shapes morphologically with an estimated diameter of 1.0 μm per shape. Surfaces were further filtered to those with volumes above 6.5943 μm^3. For the *vasa* video, surfaces were seeded with a grain size of 0.2 μm and largest sphere diameter of 2.0 μm. These were thresholded to an intensity value of 20.3128 a.u., and Imaris was instructed to split connected shapes morphologically with an estimated diameter of 1.0 μm per shape. Surfaces were further filtered to those with volumes above 5.2160 μm^3. Both sets of surfaces were artificially colored with a white-blue marbled solid coloring. Videos were curated using the “animation” feature by adding still shots and horizontal rotations respectively. Videos were assigned 500 frames divided among the shots and playback speed was set to 15 frames/sec. Additional video annotations (beyond scale bars) were made in Adobe Premiere Pro.

### Manually Quantifying Average Chromatin Intensity

DAPI-stained chromatin intensity along linescans was manually quantified in FIJI (Schindelin et al., 2012). Linescans were manually drawn along the longest axis of each nucleus. Each EIC nucleus was lined on the plane with the widest axis for that nucleus, and each EIC had a corresponding surrounding cell that was lined on that same plane. We excluded from the analyses any nuclei that appeared fixed at metaphase, evident by a bright bar of concentrated DAPI signal. Example linescans for one z plane of one embryo (2 EICs and 2 surrounding nuclei) is shown in Supp. Fig. 1B. Six embryos were analyzed, with four EICs and four surrounding cells per embryo, for 24 EICs and 24 surrounding cells in total. Linescans were measured using the “measure” function in FIJI, which reports linescan length, minimum intensity, maximum intensity, and average intensity in arbitrary units. Intensities were normalized to a manually-selected region of 1296 pixels^2 of the background, specific to each image. The average intensity in this background region was subtracted from the average intensity calculated by the measure function for each linescan. The resulting fluorescent intensities were plotted in prism for all EICs and all surrounding cells. The mean and standard deviation from the mean for the EICs and the surrounding cells were also plotted. Across the population, EIC linescan averages were lower than those of surrounding cells. An unpaired t-test was performed resulting in p<0.0001 chance that the difference seen between the two datasets was due to random chance, indicating that the DAPI-stained chromatin in the EICs had a significantly lower fluorescence intensity than that in the surrounding cells along the linescans taken.

## Supporting information

Graphical Abstract

## Figure Legends

**Supplementary Figure 1.**
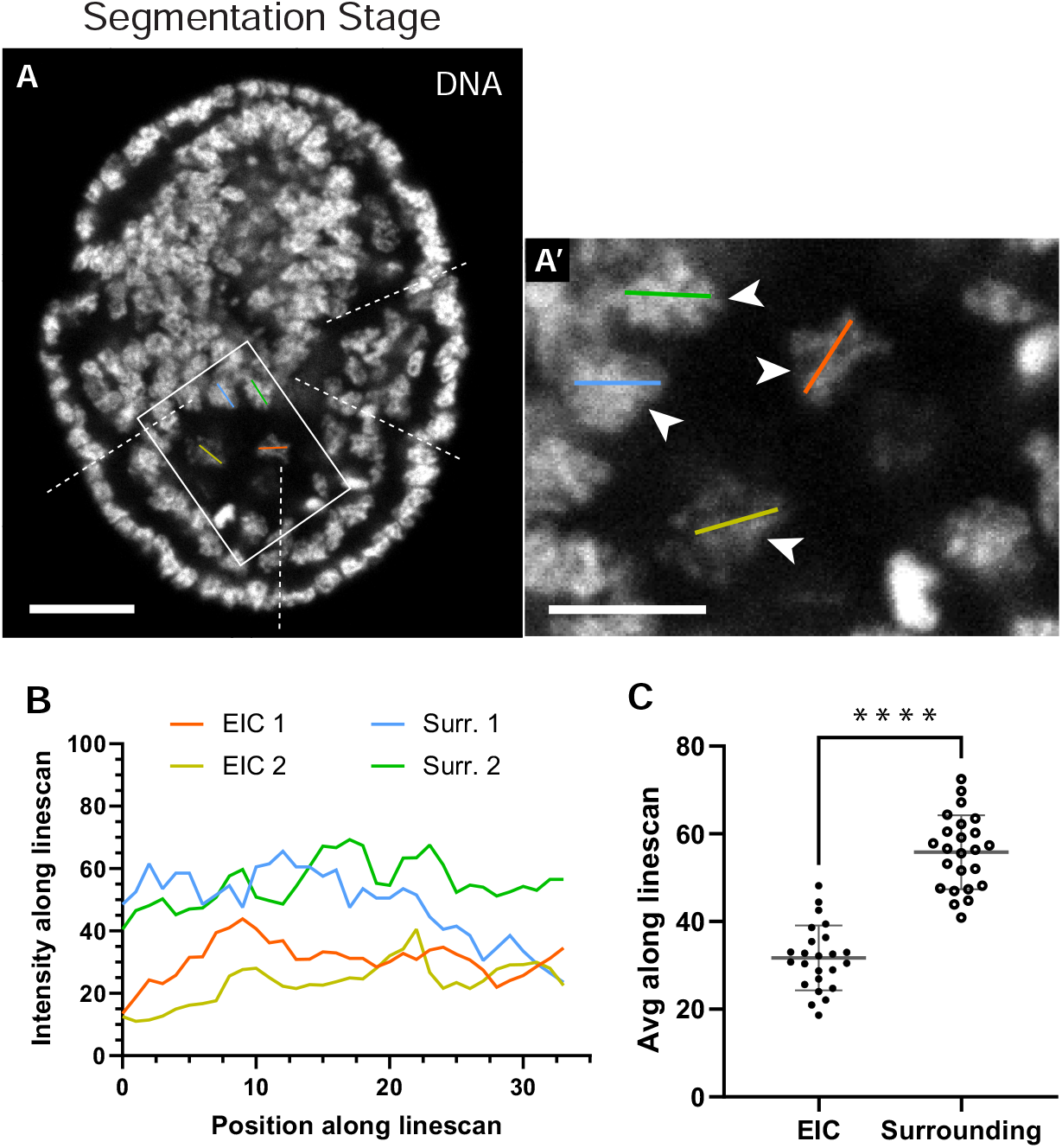
The four EICs exhibit diffuse chromatin morphology. (A) The single z plane of a DAPI-stained embryo from Fig. 1 B and further enlarged view of the boxed region indicating the positions of the linescans taken over two EIC nuclei (orange and yellow lines) and two surrounding nuclei (green and blue lines), all indicated by arrowheads in the enlarged view, (A, scale bar = 10 μm, and A’, scale bar of enlarged region = 4 μm). (B) Plotted intensities along the linescans from A. (C) Average intensity along all linescans from EIC nuclei and surrounding nuclei (p<0.0001, t-test, unpaired, n = 6 embryos, 4 EIC and 4 surrounding nuclei per embryo, bars indicate mean and standard deviation from the mean). EIC nuclei had significantly lower average DAPI signal intensity across linescans than did surrounding nuclei.

**Supplementary Figure 2.**
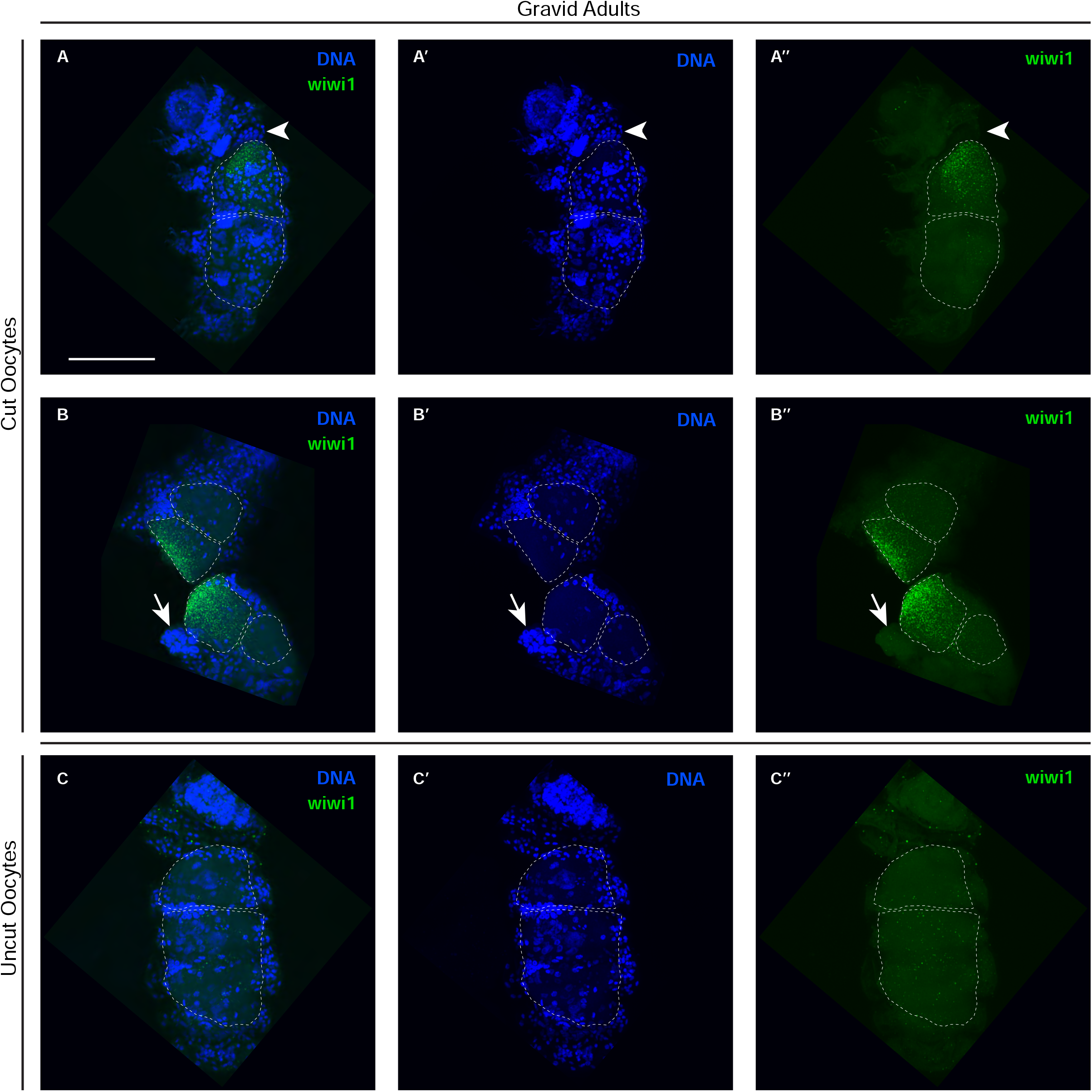
*wiwi1* mRNA is present in developing oocytes. (A and B) Presence of *wiwi1* mRNAs by FISH in oocytes in gravid adults. (C) Staining appears absent in gravid adults for which oocytes were not cut in the permeabilization process. Arrowhead indicates location of previously-identified germ stem cell niche, which appears not enriched for *wiwi1*, despite appearing cut. Arrow indicates region where presumptive somatic tissue is clearly cut but is not enriched for *wiwi1*. (n = 3 experiments, 5 (3 cut and 2 uncut oocytes), 6 (4 cut and 2 uncut oocytes), and 10 (6 cut and 4 uncut oocytes) gravid adults, respectively, scale bar = 50 μm)

**Supplementary Figure 3.**
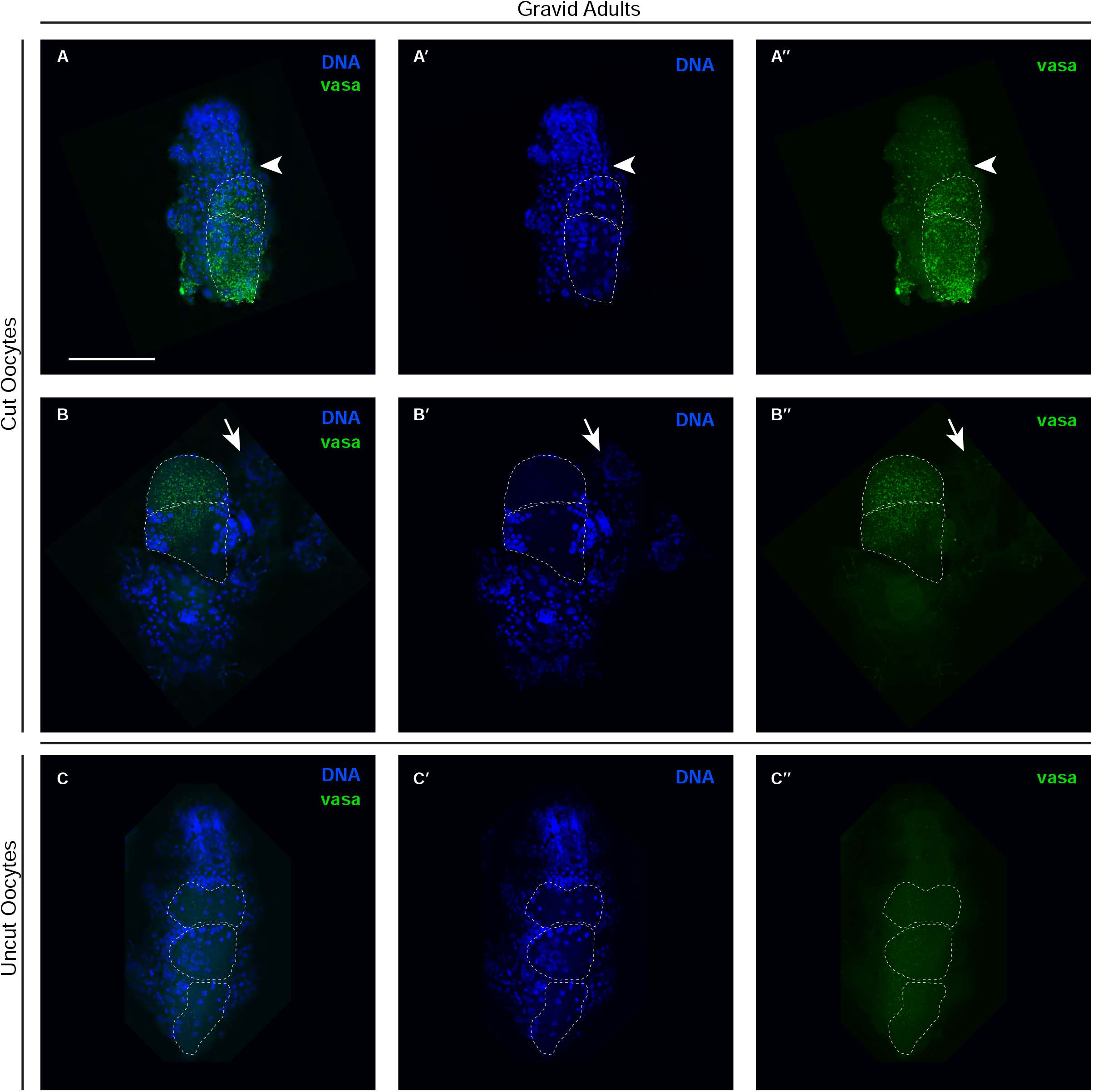
*vasa* mRNA is present in developing oocytes. (A and B) Presence of *vasa* mRNAs by FISH in oocytes in gravid adults (A and B). (C) Staining appears absent in gravid adults for which oocytes were not cut in the permeabilization process. Arrowhead indicates location of previously-identified germ stem cell niche, which appears not enriched for *vasa*, despite appearing cut. Arrow indicates region where presumptive somatic tissue is clearly cut but is not enriched for *vasa*. (n = 1 experiment, 9 gravid adults, 5 with cut oocytes and 4 with uncut oocytes, scale bar = 50 μm)

**Supplementary Figure 4.**
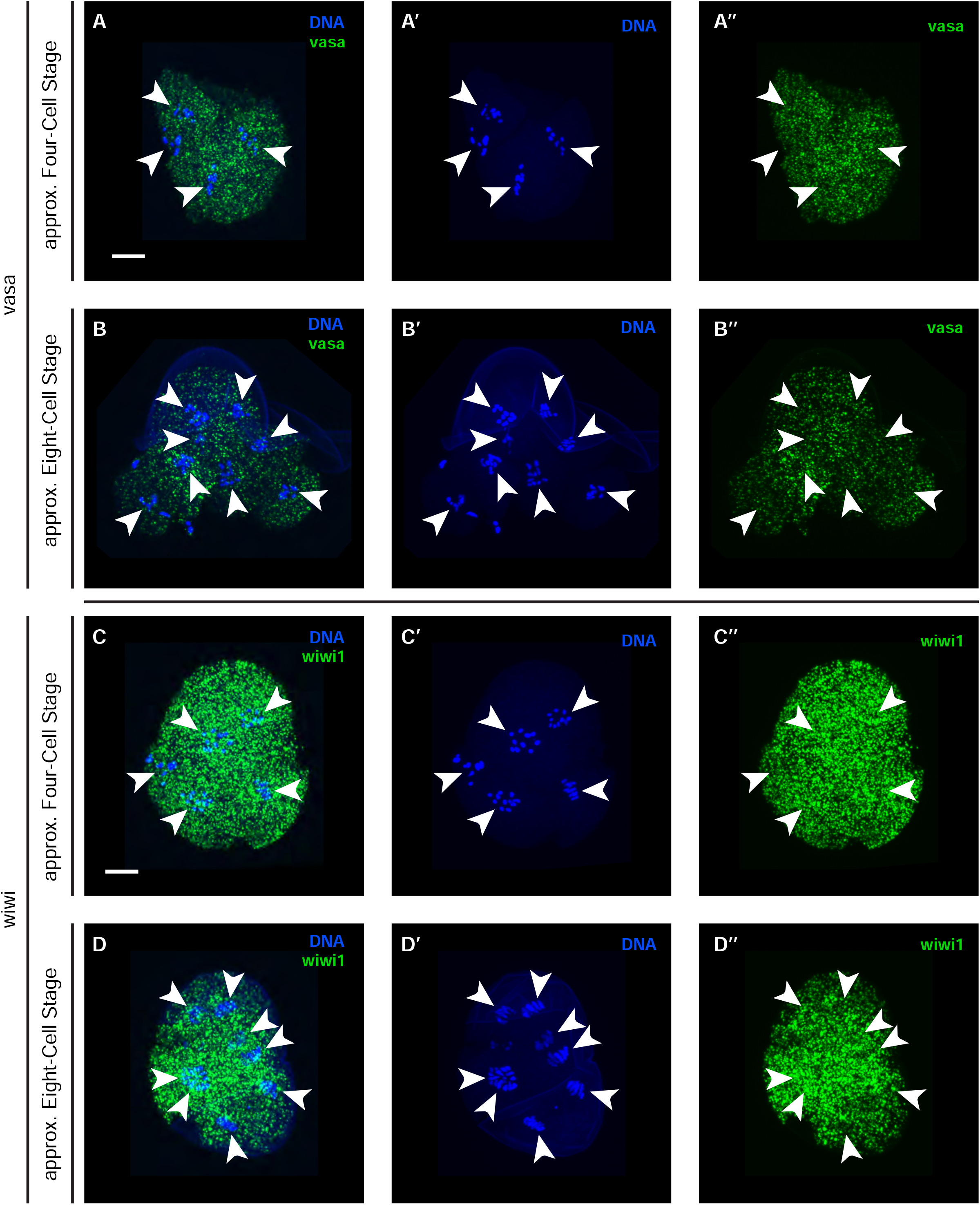
*vasa* and *wiwi1* mRNAs are uniformly distributed in early cleavage stage embryos. (A and B) Presence of *vasa* mRNAs by FISH in early cleavage stages: (A) approximately four-cell stage embryo and (B) approximately eight-cell stage embryo. (C and D) Presence of *wiwi1* mRNAs by FISH in early cleavage stages: (C) approximately four-cell stage embryo and (D) approximately eight-cell stage embryo. Arrowheads indicate nuclei. (*vasa* n = 1 experiment, 2 embryos, *wiwi1*, n = 1 experiment, 4 embryos, scale bars = 10 μm)

## Tables

Table 1. Accession numbers for phylogenetic analyses.

*For some, accession numbers for protein sequences could not be found, so the online translation tool on ExPASy was used to translate available RNA nucleotide sequences into amino acids (https://web.expasy.org/translate/). In these cases, the nucleotide sequence used matches the accession number listed in the “transcript accession” column. Translation frame and direction was confirmed by analyzing the domains of the resulting amino acid sequence in the NCBI Conserved Domain database and comparing against the domains of *D. melanogaster* homologs as a point of reference.

Table 2. Primers used for cloning.

*Modified primers for M13 fwd and rev with higher melting temperatures than the original M13 primers were developed specific to the pCR4-TOPO vector.

## Videos

Video 1. Migrating EICs

Single plane DIC microscopy video of EIC migration from after internalization from the ventral surface to the segmentation stage. Approximate positions of the four EIC nuclei marked by asterisks (nuclei evident by the absence of yolk). In some planes, all four EIC nuclei are visible, and in other planes, less than four nuclei are visible, due to the motile nature of the four EICs. Embryo is oriented with anterior to the left and posterior to the right. Embryo begins oriented with the ventral side up and obliquely towards the coverslip. The embryo rotates towards the coverslip during the epithelium stage to be ventral side down and obliquely towards the coverslip. Text annotations are included to note when the z plane is shifted to continue tracking the four EICs. Text annotations are also included to identify the embryonic stages: EIC migration after internalization, epithelium, elongation, and segmentation. Additional arrow annotations are drawn to indicate the axis of elongation during the elongation stage and the approximate positions of the four ectodermal constriction points that approximately align with the positions of segment boundaries during the segmentation stage. Time in minutes from the start to end of the video is shown in the top left corner. Each frame was taken every minute. Playback speed is 15 frames/sec. (scale bar = 10 μm)

Video 2. Rotating nuclei showing position and enrichment of *wiwi1*.

Video of *wiwi1* and DAPI-stained chromatin signal in a segmentation stage (24 hpl) embryo curated in Imaris 9.9.1. Embryo is initially oriented anterior up, posterior down, dorsal left, and ventral right. Embryo is rotated in a right-handed fashion about the anterior-posterior axis. Chromatin signal is in blue and *wiwi1* is in green. The region enriched for *wiwi1* was manually cropped, and the chromatin signal in this region is displayed to reveal that at least four nuclei appear to be surrounded by the enriched *wiwi1* signal. Because the DAPI staining of these four nuclei was so much more diffuse than in the remainder of the embryo, we had to increase the brightness of the DAPI signal in the manually cropped region in order to better see the four nuclei of the cells enriched for *wiwi1*. The chromatin signal in the manually-cropped region was then segmented in the software to identify the four nuclei located in this region. Segmented nuclei are shown as solid objects with a white-blue marbled coloring. The segmented nuclei are overlaid on the whole embryo to show their relative position in the embryo and in the region enriched for *wiwi1*. The four segmented nuclei appear to be in the same position as the four EICs. (scale bar = 10 μm)

Video 3. Rotating nuclei showing position and enrichment of *vasa*.

Video of *vasa* and DAPI-stained chromatin signal in an elongation stage (18 hpl) embryo curated in Imaris 9.9.1. Embryo is initially oriented anterior up, posterior down, dorsal left, and ventral right. Embryo is rotated in a right-handed fashion about the anterior-posterior axis. Chromatin signal is in blue and *vasa* is in green. The region enriched for *vasa* was manually cropped, and the chromatin signal in this region is displayed to reveal that at least four nuclei appear to be surrounded by the enriched *vasa* signal. Because the DAPI staining of these four nuclei was so much more diffuse than in the remainder of the embryo, we had to increase the brightness of the DAPI signal in the manually cropped region in order to better see the four nuclei of the cells enriched for *vasa*. The chromatin signal in the manually-cropped region was then segmented in the software to identify the four nuclei located in this region. Segmented nuclei are shown as solid objects with a white-blue marbled coloring. The segmented nuclei are overlaid on the whole embryo to show their relative position in the embryo and in the region enriched for *vasa*. The four segmented nuclei appear to be in the same position as the four EICs. (scale bar = 10 μm)

## Acknowledgements

We thank members of the Goldstein lab, past and present, for their generosity of time, scientific discussion, and advice, especially Pu Zhang for an unbiased confirmation of the lineage discrepancies we found, Heather Barber for the beautiful drawings of various stages in tardigrade development, Mark Slabodnick and Ariel Pani for experimental advice, the Goldstein lab team tardigrade for their unfailing support, and Courtney Clark-Hachtel, Pu Zhang, Jon Hibshman, Emily Bowie, Addie Coke, and Taylor Medwig-Kinney for comments on the manuscript. We thank Elena Keeling for additional comments on the manuscript. We thank the generous biology community at UNC-Chapel Hill, particularly Tony Perdue and Pablo Ariel for microscopy assistance, Kale Hartmann and Jeff Sekelsky for fly embryos and assistance with staining, Isai Salas Gonzalez for assistance with phylogenetic tree building, and Elizabeth Sarkel for translation of German literature. We also thank the germ cell biology community at large, particularly Gee-Way Lin, Chun-Che Chang, Hong Han, and Paul Lasko for generously sharing Vasa antibodies. We thank Florian Jug and Mangal Prakash for introduction to and assistance with the Mastodon plugin. We thank the reviewers for their thorough and constructive feedback. This work would not be possible without the help of all the individuals listed here, and many more not listed by name.

