## Supplementary figures and images for "The Embryonic Origin of Primordial Germ Cells in the Tardigrade *Hypsibius exemplaris*"

### Graphical Abstract

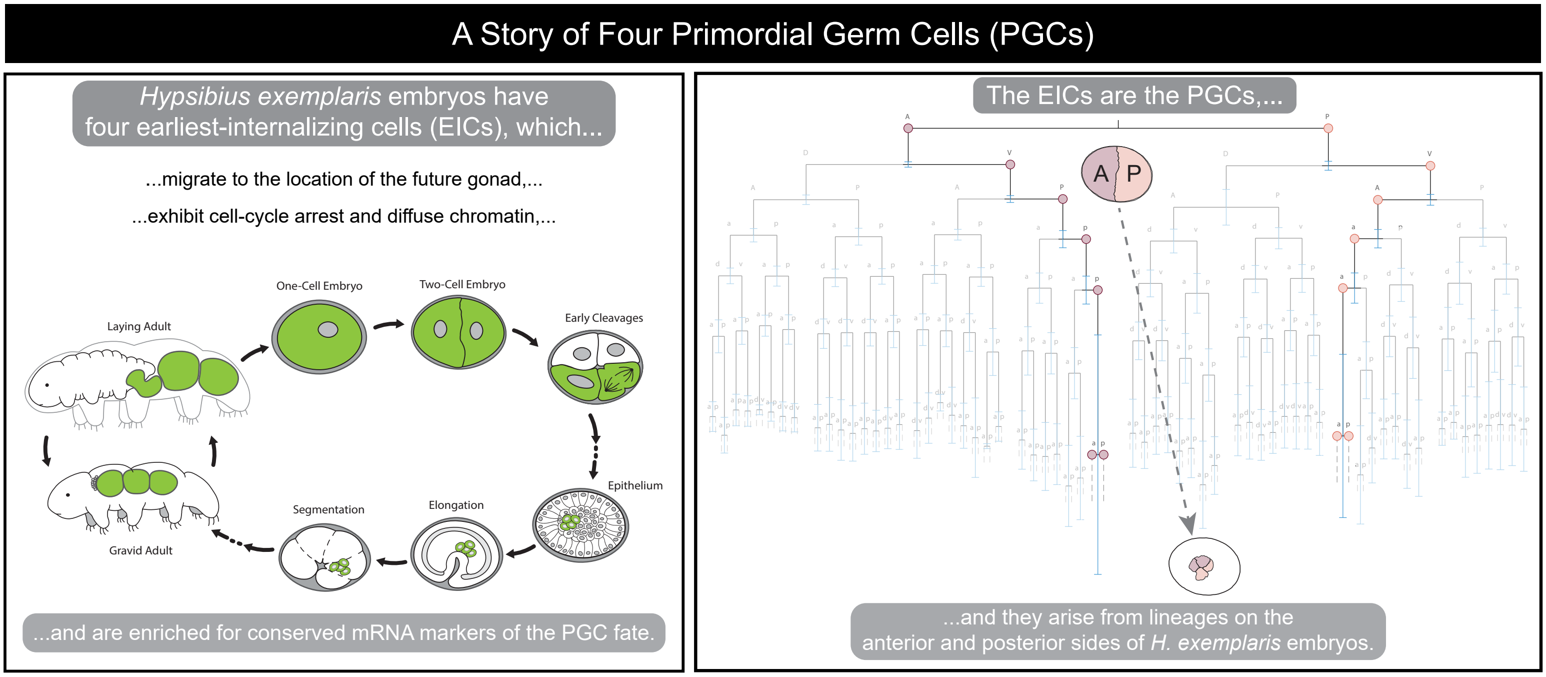


Graphical Abstract
