## Supplemental Methods for "The Embryonic Origin of Primordial Germ Cells in the Tardigrade *Hypsibius exemplaris*"

Table 1. Accession numbers for phylogenetic analyses.

| | | | | |
|---|---|---|---|---|
| **Accession Numbers** | | | | |
| **Argonautes** | | | | |
| **organism** | **species abbreviation** | **protein name** | **transcript accession** | **protein accession*** |
| *Hypsibius exemplaris* | He | He_Wiwi1 | He_tr_09087 | (expasy translation) |
| *Hypsibius exemplaris* | He | He_Wiwi2 | He_tr_08231 | (expasy translation) |
| *Hypsibius exemplaris* | He | He_AGO1 | He_tr_06607 | (expasy translation) |
| *Hypsibius exemplaris* | He | He_AGO2 | He_tr_12470 | (expasy translation) |
| *Hypsibius exemplaris* | He | He_AGO3 | He_tr_08096 | (expasy translation) |
| *Drosophila melanogaster* | Dm | Dm_Piwi | Dmel_CG6122 | NP_476875.1 |
| *Drosophila melanogaster* | Dm | Dm_Aubergine | Dmel_CG6137 | NP_476734.1 |
| *Drosophila melanogaster* | Dm | Dm_Ago3 | Dmel_CG40300 | NP_001036629.2 |
| *Caenorhabditis elegans* | Ce | Ce_PRG-1 | CELE_D2030.6 | NP_492121.1 |
| *Danio rerio* | Dr | Dr_ZIWI | NM_183338.1 | NP_899181.1 |
| *Danio rerio* | Dr | Dr_ZILI | NM_001365624.1 | NP_001352553.1 |
| *Mus musculus* | Mm | Mm_MIWI | NM_021311.3 | NP_067286.1 |
| *Mus musculus* | Mm | Mm_MIWI2 | NM_001368831.1 | NP_001355760.1 |
| *Mus musculus* | Mm | Mm_MILI | NM_001364321.1 | NP_001351250.1 |
| *Platynereis dumerilii* | Pd | Pd_Piwi | AM076487.1 | AM076487 |
| *Crassostrea gigas* | Cg | Cg_Piwi | XM_034482535.1 | LOC105339049 |
| **RNA Helicases** | | | | |
| **organism** | **species abbreviation** | **protein name** | **transcript accession** | **protein accession*** |
| *Hypsibius exemplaris* | He | He_Vasa | He_tr_12746 | (expasy translation) |
| *Hypsibius exemplaris* | He | He_Belle | He_tr_00485 | (expasy translation) |
| *Drosophila melanogaster* | Dm | Dm_Vasa | FBgn0283442 | NP723899 |
| *Drosophila melanogaster* | Dm | Dm_Belle | FBgn0263231 | NP536783 |

| | | | | |
|---|---|---|---|---|
| *Caenorhabditis elegans* | Ce | Ce_GLH-1 | WBGene00001598 | NP_491963.1 |
| *Caenorhabditis elegans* | Ce | Ce_GLH-4 | WBGene00001601 | NP_491207.3 |
| *Gryllus bimaculatus* | Gb | Gb_Vasa | AB378065.1 | BAG65665.1 |
| *Tribolium castaneum* | Tc | Tc_Vasa | NM_001039431.2 | NP001034520 |
| *Parhyale hawaiensis* | Ph | Ph_Vasa | EU289291.1 | ABX76969.1 |
| *Danio rerio* | Dr | Dr_DDX4 | ZDB-GENE-990415-272 | XP_005156510.1 |
| *Danio rerio* | Dr | Dr_PL10 | ZDB-GENE-980526-150 | XP_005168845.1 |
| *Xenopus laevis* | Xl | Xl_VLG1 | XB-GENE-1016814 | NP001081728 |
| *Xenopus laevis* | Xl | Xl_Pl10 | XB-GENE-999738 | NP001080283 |
| *Mus musculus* | Mm | Mm_Mvls | MGI:102670 | XP_011242921.1 |
| *Mus musculus* | Mm | Mm_PL10 | MGI:91842 | NP149068 |
| *Schistocerca gregaria* | Sg | Sg_Vasa | AF510054.1 | AAO15914 |
| *Nasonia vitripennis* | Nv | Nv_Vasa | NV16961 | XP_001603956.3 |
| *Apis mellifera* | Am | Am_Vasa | GB42306 | XP_006571765.2 |
| *Urechis unicinctus* | Uu | Uu_Vasa | JQ665715.1 | AFG17059.1 |
| *Platynereis dumerilii* | Pd | Pd_Vasa | AM114778.1 | CAJ38803.1 |
| *Crassostrea gigas* | Cg | Cg_Vasa | AY423380.1 | AAR37337.1 |
| *Homo sapiens* | Hs | Hs_Vasa | NM_024415.3 | NP077726 |

*For some, accession numbers for protein sequences could not be found, so the online translation tool on ExPASy was used to translate available RNA nucleotide sequences into amino acids (https://web.expasy.org/translate/). In these cases, the nucleotide sequence used matches the accession number listed in the “transcript accession” column. Translation frame and direction was confirmed by analyzing the domains of the resulting amino acid sequence in the NCBI Conserved Domain database and comparing against the domains of *D. melanogaster* homologs as a point of reference.

Table 2. Primers used for cloning.

| **Primers** | | |
|---|---|---|
| **gene** | **direction** | **sequence** |
| *wiwi1* | F1 | 5'-ATCTCTAGACCCGCGCAATG-3' |
| | F2 | 5'-GCGTTTCCGAATCATCAGGC-3' |
| | R1 | 5'-TCTTCTGCAGCATCACCTGG-3' |
| | R2 | 5'-CCTGGGTGATGCAGGGATTT-3' |
| *vasa* | F1 | 5'-CCGCACACTAGAGAGGCAAA-3' |
| | F2 | 5'-AAACCGGTCAGAAAAGTGGC-3' |
| | R1 | 5'-GAGGGTCCAAATCCACCAGG-3' |
| | R2 | 5'-TACGTCCTGTCTGTGGGAGG-3' |
| *M13 Modified | F | 5'-GTAAAACGACGGCCAGTGAATTGTAAT-3' |
| | R | 5'-CAGGAAACAGCTATGACCATGATTACG-3' |

*Modified primers for M13 fwd and rev with higher melting temperatures than the original M13 primers were developed specific to the pCR4-TOPO vector.

### Supplemental Methods

FLUORESCENCE *IN SITU* HYBRIDIZATION IN *HYPSIBIUS EXEMPLARIS* BY TYRAMIDE SIGNAL AMPLIFICATION

This protocol is modified from:

Smith, F.W. 2018. Embryonic In Situ Hybridization for the Tardigrade *Hypsibius exemplaris. Cold Spring Harbor Protocols* 2018 (11), pdb. prot102350.

#### MATERIALS

##### Reagents

Acetic anhydride

Anti-Digoxigenin-POD, Fab fragments (anti-Dig-POD) (Roche 11207733910)

AP developing solution

BM Purple AP substrate solution (Roche 11442074001)

Chymotrypsin/chitinase solution

DIG-labeled riboprobe

Fixative solution for *Hypsibius* in situ hybridization

Hybridization buffer (hyb buffer)

Plain hybridization buffer (plain hyb buffer)

Maleic acid buffer (MAB)

Methanol (25%, 50%, 70%, and 90% [v/v] in 0.5× PBTw; 100%)

Mounting medium (e.g., Fluoromount-G [Invitrogen 00495802])

PBTween (PBTw; 0.5×)

Salmon sperm DNA solution (Invitrogen 15632011), diluted in H2O to a concentration of 100 µg/mL

Spring water

SSC plus 0.1% CHAPS/Tween

Triethanolamine (1% in 0.5× PBTw)
TSA Plus Cyanine 3 50-150 slides (Akoya Biosciences NEL744001KT)

**Supplies**

Boekel Rocker II

Collection tubes (2 mL)

Compound microscope using DIC optics or laser scanning confocal microscope Depression slides

Dissecting microscope

Heat block at 100ºC

Hybridization oven at 60ºC

Microcentrifuge

Microcentrifuge tubes (1.5 mL, low-retention)

Mobicols (Boca Scientific)

Mobicol mini-columns with Luer-lock cap, closing cap, and plug (M1002)

Large filters for Mobicols, 10-µm pore size (M2210)

Bottom filters for Mobicols, 10-µm pore size (M2110)

Needles (25 gauge)

Pasteur pipettes (9 inch, glass) Petri dishes (35 mm × 10 mm) Syringes (1 mL)

U-bottom culture plate (96 well)

Vortexer (VWR Genie 2 G560)

Water bath at 60ºC

**METHOD**

*When embryos are in 1.5 ml tubes, wash steps include a centrifugation step at 3000 rcf for 3 minutes. Throughout the protocol, make sure that the embryos remain submerged in liquid. All wash steps are at room temperature unless otherwise indicated.*

**Embryo Collection**

1. In a 35 mm × 10 mm Petri dish filled with spring water, slice through the middle of maternal exuviae filled with embryos with a 25-gauge needle attached to a sterile 1-mL syringe.

2. Transfer embryos to a 1.5-mL tube filled with 0.5× PBTw.

**Primary Permeabilization**

3. Centrifuge the 1.5-mL tube containing embryos at 18,500g for 3 min. Remove most of the liquid, leaving embryos suspended in ~20 µL of 0.5× PBTw.

4. Add 20 µL of chymotrypsin/chitinase solution, and let stand for 1 h. Centrifuge at 3000g for 3 min, and remove most of the liquid.

5. Wash three times as follows: Add 500 µL of 0.5×PBTw, let stand for 5 min, centrifuge at 3000g for 3 min, and remove most of the liquid.

**Paraformaldehyde Fixation**

6. Wash with 1 mL of fixative solution while shaking vigorously on a VWR Genie 2 G560 Vortexer set to shake speed 3 for 30 min. Centrifuge at 3000g for 3 min, and remove most of the liquid.

7. Wash five times as follows: Add 500 µL of 0.5×PBTw, let stand for 5 min, centrifuge at 3000g for 3 min, and remove most of the liquid.

**Cleaning Mobicol Mini-Columns**

8. Follow the manufacturer's instructions to set up mini-columns:

i. Unscrew the Leur-lock cap, add 600 µL of 0.5× PBTw to the top of the mini-column, and then screw the Leur-lock cap and syringe back on.

ii. Push the syringe plunger all the way down, and discard the liquid that collects in the tube.

**Methanol Dehydration**

*For all mini-column washes below, unscrew the Leur-lock cap, add the liquid to be used for washing the embryos, and then screw the Leur-lock cap and syringe back on. After letting the embryos stand in the liquid for the specified time, push the liquid through the mini-column with a 1-mL syringe. The embryos must remain submerged in liquid at all times, so during these washes, pull back on the syringe plunger to form a vacuum in the mini-column to ensure that not all of the liquid is lost.*

9. Transfer embryos to a clean mini-column using a 9" glass Pasteur pipette.

10. Wash embryos for 5 min in 500 µL of 25% methanol in 0.5× PBTw.

11. Repeat Step 10 with 50%, 70%, and 90% methanol in 0.5× PBTw.

12. Quickly wash the embryos three times in 500 µL of 100% methanol. After adding the final wash of 100% methanol, insert a bottom plug into the column and replace the Luer-lock cap with a closing cap. Leave the tube at −20ºC for at least 20 min.

**Methanol Rehydration**

13. Remove the bottom plug, and replace the closing cap with the Luer-lock cap and syringe. Wash the embryos for 5 min each in 500 µL of 90%, 70%, 50%, and 25% methanol in 0.5× PBTw, followed by three quick washes with 500 µL 0.5× PBTw.

**Secondary Permeabilization**

14. Use a 9 inch glass Pasteur pipette to transfer the embryos from the mini-column into a 35-mm dish filled halfway with 0.5× PBTw.

15. Cut the embryos out of the eggshells with a 25-gauge needle attached to a 1-mL syringe.

16. Re-collect the embryos in a clean mini-column in 500 µL of 0.5×PBTw, and then push the excess liquid through the mini-column with a 1-mL syringe.

**Acetylation**

17. Quickly wash the embryos with 500 µL of 0.5× PBTw.

18. Quickly wash the embryos twice with 500 µL of 1% triethanolamine in 0.5× PBTw.

19. In a 1.5-mL tube, combine 1.3 µL of acetic anhydride and 500 µL of 1% triethanolamine in 0.5× PBTw, and vortex briefly. Wash the embryos for 5 min in this solution.

20. Repeat step 19.

21. Quickly wash the embryos twice with 500 µL of 0.5× PBTw.

**Prehybridization**

22. Repeat Step 21 one more time, but at the end of the wash, pass half of the 0.5×PBTw through the mini-column and replace it with room-temperature hyb buffer. Rock gently on a Boekel Rocker II for 20 min at room temperature.

*The syringe remains attached during the washes in Steps 22–24. Ensure that the embryos remain submerged in liquid at all times by pulling back on the syringe plunger to form a vacuum in the mini-column.*

23. Wash with 500 µL of room-temperature hyb buffer, rocking gently for 20 min at room temperature.

24. Wash with 500 µL of hyb buffer heated to 60ºC in a water bath. Let the embryos stand in a hybridization oven for 2 h at 60ºC.

**Hybridization**

25. Add DIG-labeled riboprobe to a final concentration of ~0.5 µg/mL in 500 µL of hyb buffer. Incubate for 5 min at 100ºC on a heat block.

26. Boil salmon sperm DNA, and shear by pulling in and out of a 25-gauge needle attached to a 1-mL syringe. Add 1.0 µL of 100 µg/mL boiled and sheared salmon sperm DNA to the DIG-labeled riboprobe solution.

27. Pass most of the buffer that the embryos are suspended in through the mini-column, and add the riboprobe solution.

28. Hybridize overnight at 60°C in a hybridization oven.

*The syringe continues to remain attached during Steps 28–32. Ensure that the embryos remain submerged in liquid at all times by pulling back on the syringe plunger to form a vacuum in the mini-column.*

**Post-hybridization Washes**

*For post-hybridization washes, all buffers should be preheated to 60°C and the embryos should be incubated in a 60°C hybridization oven.*

29. Quickly wash the embryos ten times with 500 µL of plain hyb buffer.

30. Quickly wash the embryos six times with 500 µL of 2× SSC plus 0.1% CHAPS/Tween.

31. Quickly wash the embryos twice with 500 µL of 1× SSC plus 0.1% CHAPS/Tween.

32. Quickly wash the embryos twice with 500 µL of 0.2× SSC plus 0.1% CHAPS/Tween.

**Immunohistochemistry**

33. Remove the embryos from the hybridization oven. Quickly wash the embryos twice with 500 µL of room-temperature 0.5× PBTw.

34. Incubate the embryos for 2 h at room temperature in 500 µL of the blocking buffer from the TSA kit (Akoya Biosciences).

35. Incubate the embryos overnight at 4ºC in 500 µL of 1:1500 anti-DIG-POD antibody:blocking buffer.

**Post-antibody Washes**

36. Quickly wash the embryos ten times with 500 µl MAB buffer.

**Tyramide Signal Amplification**

37. Wash the embryos with 500 µl of 1X amplification diluent from the TSA kit (Akoya Biosciences). Rock gently for 5 min.

38. Prepare 200 µL of TSA Plus working solution (4 µL Cyanine 3 Plus Amplification Reagent: 196 µL 1X amplification diluent from TSA kit [Akoya Biosciences]).

39. Wash the embryos with 200 µL of TSA Plus working solution. Rock gently for 30 min in the dark.

40. Preheat 0.5× PBTw to 60ºC.

41. Quickly wash the embryos five times with 0.5× PBTw preheated to 60ºC.

42. Wash the embryos five times for 10 min in 0.5× PBTw preheated to 60ºC while incubating at 60ºC.

*The embryos can be stored at 4ºC in 0.5× PBTw until they are mounted.*

**Imaging**

43. Transfer embryos to a clean depression slide using a 9” glass Pasteur pipette.

44. Transfer all of the embryos to one well of a 96-well plate filled with 200 µL of 0.5× PBTw.

45. Mount specimens on microscope slides in Fluoromount-G (Invitrogen) or other appropriate mounting medium.

46. Image on laser scanning confocal microscope.

### RECIPES

Chymotrypsin/chitinase solution

| | |
|---|---|
| Chitinase (Sigma-Aldrich C6137) | 5 units |
| Chymotrypsin (Sigma-Aldrich C4129) | 10 mg |
| 0.5× PBS (diluted from 10× PBS [pH 7.4] | 1 m |
| Final volume | ~1 ml |

50× Denhardt’s

| | |
|---|---|
| Ficoll | 5 g |
| polyvinylpyrrolidone | 5 g |
| Bovine Serum Albumin | 5 g |
| $ddH_2O$ | to 500 ml |
| Final volume | 500 ml |

Fixative Solution

| | |
|---|---|
| Heptane | 333.33 µl |
| 16 % formaldehyde (Electron Microscopy Sciences 15700) | 250 µl |
| 10% Tween20 | 10 µl |
| 0.5× PBTw | 406.67 µl |
| Final volume | ~1 ml |

Hybridization buffer

| | |
|---|---|
| 100% Formamide | 25 ml |
| 20X SSC (pH 7.0) | 12.5 ml |
| 100 mg/ml heparin | 50 µl |
| 10% Tween20 | 500 µl |
| 10 mg/ml yeast RNA | 500 µl |
| 10% CHAPS | 500 µl |
| 50X Denhardt’s | 1 ml |
| DEPC $H_2O$ | 9.95 ml |
| Final volume | 50 ml |

Maleic Acid Buffer

| | |
|---|---|
| Maleic Acid | 11.61 g |
| NaCl | 8.77 g |
| NaOH pellets | 7.2 g |
| Sterile $H_2O$ | 500 ml |
| adjust pH to 7.5 | |
| Fill to 1 L with $H_2O$ | |

Phosphate-Buffered Saline (PBS; 10×, pH 7.4)

| | |
|---|---|
| $Na_2HPO_4 \cdot 7H_2O$ | 25.6 g |
| NaCl | 80 g |
| KCL | 2 g |
| $KH_2PO_4$ | 2 g |
| $H_2O$ | 1 L |
| Adjust to pH 7.4. | |
| Mix and autoclave. | |
| Final volume | ~1 L |

PBTween (PBTw; 0.5×)

| | |
|---|---|
| 10X PBS | 2.5 ml |
| 10% Tween20 | 500 µl |
| DEPC $H_2O$ | 47 ml |
| Final volume | 50 ml |

Plain Hybridization Buffer (Plain Hyb Buffer)

| | |
|---|---|
| 100% Formamide | 25 ml |
| 20X SSC (pH 7.0) | 12.5 ml |
| 10% Tween20 | 500 µl |
| DEPC $H_2O$ | 12 ml |
| Final volume | 50 ml |

20X SSC buffer

| | |
|---|---|
| NaCl | 175.3 g |
| Sodium Citrate | 88.2 g |
| dd$H_2O$ | 800 ml |
| adjust pH to 7.0 | |

Fill to 1 L with dd$H_2O$. Sterilize by autoclave. Use dd$H_2O$ to dilute to working concentrations. Add CHAPS/Tween to 0.1%.
